# Genome instability footprint under rapamycin and hydroxyurea treatments

**DOI:** 10.1101/2023.02.28.530484

**Authors:** Jing Li, Simon Stenberg, Jia-Xing Yue, Ekaterina Mikhalev, Dawn Thompson, Jonas Warringer, Gianni Liti

## Abstract

The mutational processes dictating the accumulation of mutations in genomes are shaped by genetic background, environment and their interactions. Accurate quantification of mutation rates and spectra under drugs has important implications in disease treatment. Here, we used whole-genome sequencing and time-resolved growth phenotyping of yeast mutation accumulation lines to give a detailed view of the mutagenic effects of rapamycin and hydroxyurea on the genome and cell growth. Mutation rates depended on the genetic backgrounds, but were only marginally affected by rapamycin. As a remarkable exception, rapamycin treatment was associated with frequent chromosome XII amplifications, which compensated for rapamycin induced rDNA repeat contraction on this chromosome and served to maintain rDNA content homeostasis and fitness. In hydroxyurea, a wide range of mutation rates were elevated regardless of the genetic backgrounds, with a particularly high occurrence of aneuploidy that associated with dramatic fitness loss. Hydroxyurea also induced a high T-to-G transversion rate that reversed the common G/C-to-A/T bias in yeast and gave rise to a broad range of structural variants, including mtDNA deletions. The hydroxyurea mutation footprint was consistent with the activation of error-prone DNA polymerase activities and non-homologues end joining repair pathways. Taken together, our study provides an in-depth view of mutation rates and signatures in rapamycin and hydroxyurea and their impact on cell fitness, which brings insights for assessing their chronic effects on genome integrity.

## Introduction

Genome instability, referring to the accumulation of mutations, is typified by specific mutational signatures^1^. Such signatures can be shaped by external factors, such as drug treatments, that target different aspects of the cellular machinery. Understanding why particular mutations occur and how they vary between individuals or environments is fundamental to widen our knowledge of the underlying mechanisms of mutagenesis^2,3^. However, accurate estimation of mutation rates and profiling of mutational signatures in specific environments are challenging to achieve. This is partially due to the confounding effects of selection, which favors some variants over others and thereby affects their frequency and likelihood of detection. Additionally, a continuous spatio-temporal record of cell populations is hard to obtain, which makes it difficult to track the alteration of genetic and phenotypic properties during evolution. Therefore, it is crucial to examine mutational signatures and their phenotypic consequences under conditions where selection can be minimized, and in a time-resolved manner.

The budding yeast *Saccharomyces cerevisiae* is a single-celled, eukaryotic model organism that can be easily manipulated and controlled in the lab. Asexually reproducing populations of yeast can be propagated in the lab for thousands of generations as Mutation Accumulation Lines (MALs). This experimental evolution protocol minimizes selection by repeatedly forcing cell populations through bottlenecks of random, single cells and mutations are therefore accumulated in a largely unbiased way. Intermediate stages of MALs evolution can be stored in freezers, ensuring that a fossil record of the evolving cells is preserved. The coupling of MALs with the usage of highly parallelized cell growth phenotyping platforms and whole-genome-sequencing allows us to monitor the fitness trajectories of the lines and to reconduct them to genomic alterations accumulating over time^4–8^. Yeast MALs have been used to estimate mutation rates and spectra for common lab strains in rich medium^9,10^, as well as the effects of controlled variation in ploidy^11–14^ and of some specific disruptions in the DNA repair/replication machinery^15,16^. However, mutational patterns can be also shaped by environmental factors, such as drugs and diet^17–21^, and may differ between genetic backgrounds^22^. So far, systematic studies that explore the mutational landscape under stress conditions, particularly long-term anticancer treatments, are lacking.

Here, we compared the mutation profiles of 96 mutation accumulation lines (MALs), derived from 10 distinct genetic backgrounds, that were evolved for a total of ~181,440 generations in a drug-free (rich medium; YPD) condition, stressful hydroxyurea (HU) and rapamycin (RM) conditions. HU and RM represent two distinct mechanisms of action of chemotherapeutic treatments: impairing DNA repair/replication and inhibiting the TOR signaling in proliferating cells, respectively. The mutation rates and spectra of these two drugs were highly distinct at both nucleotide and structural levels, and this had direct effects on cell fitness. Our study represents an effective approach for the assessment of mutagenic risks of drugs and the understanding of their long-term effects.

## Results

### Mutation accumulation under HU but not RM impairs cell growth

To avoid confounding effects from working with a single and potentially atypical genetic background, we selected ten diploid *S. cerevisiae* genetic backgrounds and propagated MALs in hydroxyurea (HU) 10 mg/ml, rapamycin (RM) 0.025 μg/ml and drug-free rich medium (YPD, as control) (Fig. 1a, Table S1). The 10 genetic backgrounds comprised four homozygous representatives of diverged *S. cerevisiae* lineages^23^ (WA: West African, WE: Wine/European, NA: North American, SA: Sake), as well as six heterozygous hybrids obtained through pairwise crosses of their haploid offspring (WA/WE, NA/WE, NA/WA, SA/WE, SA/WA, SA/NA). The growth of these 10 genetic backgrounds varied under drug exposures (Fig. 1b, Tables S2-S4). The mean cell doubling time increased by 43.2% (standard deviation (sd) across strains=5.6%) and 38.1% (sd=16.0%) in HU and RM respectively compared with drug-free medium. Correspondingly, the mean absolute cell yield declined by 70.3% (sd=11.01%) in HU and 77.3% (sd=4.7%) in RM and the mean cell viability was 77.2% (sd=21.5%) and 94.7% (sd=7.9%) in HU and RM relative to in YPD.

**Figure 1.**
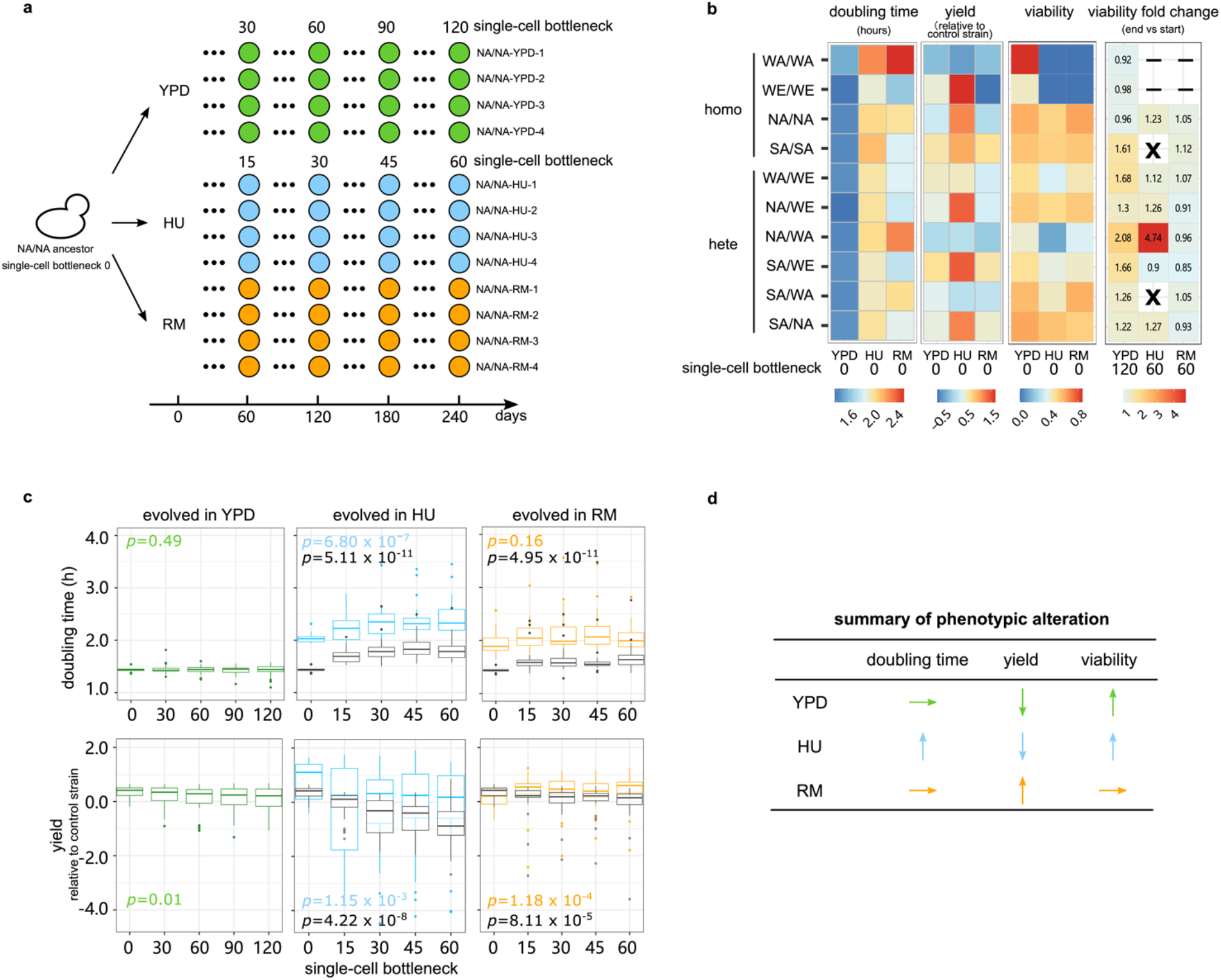
MALs design and fitness. **(a)** We evolved MALs for a total of 120 (YPD) and 60 (HU and RM) single-cell bottlenecks. The ten genetic backgrounds included four diploid homozygous strains (WA - West African, WE - Wine/European, NA - North American, SA - Sake) and all six possible hybrids derived from them. For each strain, we initiated 4 replicated MALs in each condition named as “strain-condition–replicate”, with e.g., “NA/NA-YPD-1” denoting the North American homozygote, evolved in YPD, replicate 1. Green, blue and orange circles represent the MALs in YPD, HU and RM respectively. The NA/NA genetic background MALs overview is shown as example. **(b)** Heatmap show the cell doubling time (hours), cell yield (log2 cell yield normalized to that of the spatial control strain, NA/WA) and cell viability. Cell viability in YPD is represented as CFU^YPD^ counted cells, while cell viability in HU and RM is calculated as CFU^HU or RM^/CFU^YPD^). “X” indicates MALs with all four replicates going extinct. “-” indicates strains excluded because of extremely low fitness in that condition and could not be propagated. **(c)** Evolution of cell doubling time and yield. Boxplots show the cell doubling time and yield in drug and drug-free conditions (see legend) across the single-cell bottleneck (x-axis). Center line: median; box: interquartile range (IQR); whiskers: 1.5 ×IQR; dots: outliers beyond 1.5 ×IQR. The *p* value is measured by Mann–Whitney U test by comparing phenotypes at the initial and last timepoints of evolution. For each condition, the number of MALs included in the analysis (*N*) is *N_YPD_*=40, *N_HU_*=24, *N_RM_*=32. **(d)** Qualitative summary of the direction in which each fitness proxy changes during the MALs.

We evolved four replicates of each of the 10 genetic backgrounds over 120 single-cell bottlenecks in YPD and 60 single-cell bottlenecks in HU and RM, corresponding to approximately 2760, 1200 and 1320 consecutive cell divisions respectively. Hereafter, we use the nomenclature “strain–condition-replicate” to represent each MAL (Fig 1a). Cells of two homozygous genetic backgrounds (WA/WA, WE/WE) showed too low viability in HU and RM for the MAL evolution to be initiated and another two (SA/SA and SA/WA) went repeatedly extinct across all replicates during the MAL evolution in HU (Fig. 1b). Therefore, a total of 40, 24 and 32 MALs survived to the end of the MAL evolution in YPD, HU and RM respectively (Table S1). We stored all the MALs at five intermediate stages of evolution to generate frozen fossil records (Fig. 1a). For each of the 96 surviving MALs, we established a dense cell growth record by measuring the cell doubling time and the normalized cell yield as proxies for fitness over the course of evolution (Fig. 1c, Supplementary Fig. 1 and 2, Table S2). In YPD, the cell doubling time remained stable for all the MALs (mean 1.44 h initial *vs*. 1.43 h last time points), while the cell yield declined somewhat (mean log_2_ normalized yield 0.35 initial *vs*. 0.07 last time points, *p* = 0.01). The cell yield decline was largely due to four lines becoming *petite*, i.e. forming very small colonies, which is a signature of respiration deficiency, and one line becoming haploid (Supplementary Fig. 2a).

**Figure 2.**
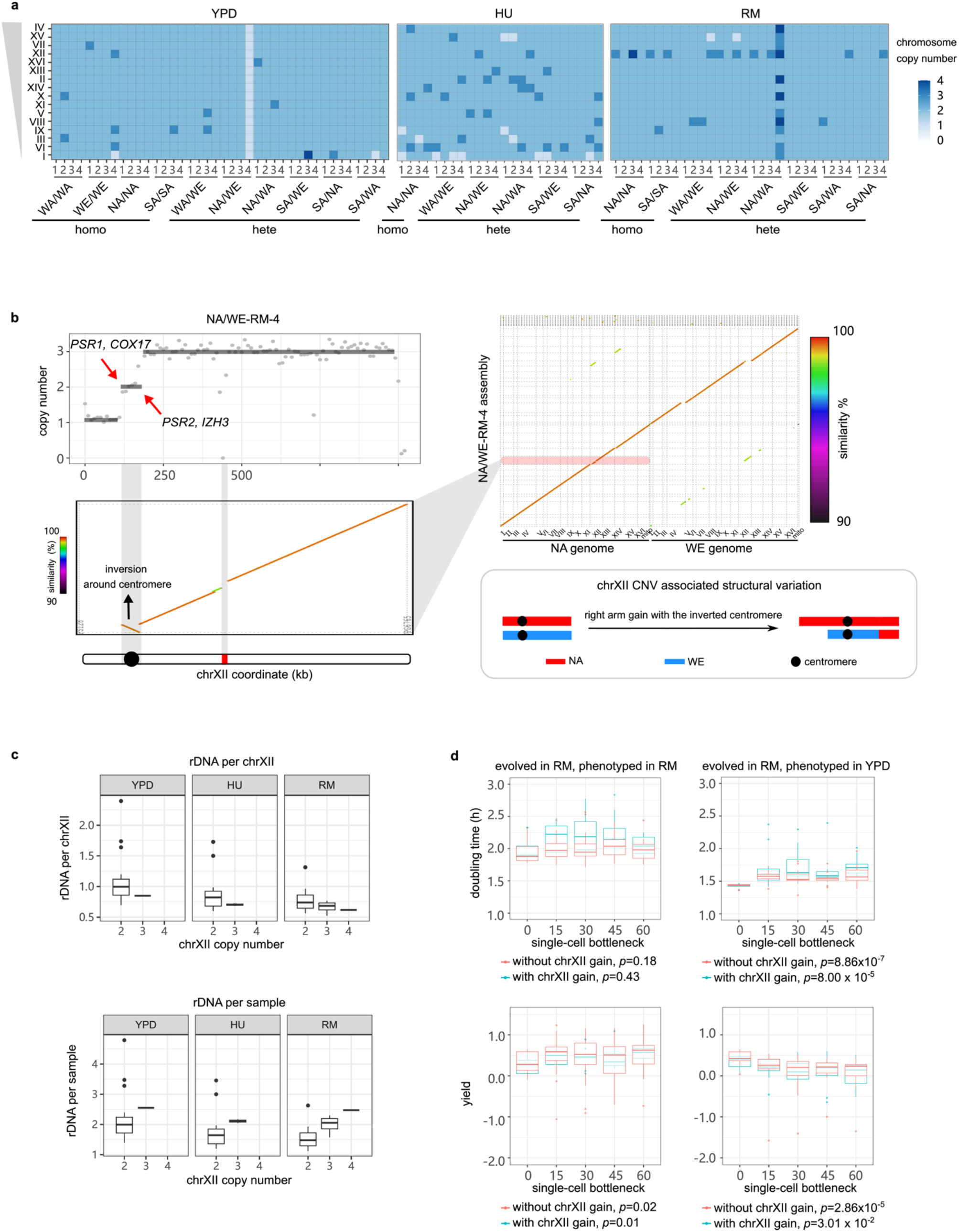
Chromosome XII amplification compensates for rDNA contraction in RM. **(a)** Aneuploidies identified in the MALs. Blue tones indicate chromosome copy number. The chromosomes are sorted by their size with larger chromosomes on the top. **(b)** Upper left, the short read sequencing revealed segmental CNV on chromosome XII of NA/WE-RM-4. The red arrows indicate the breakpoint of each segmental CNV. Upper right dot plot, the x-axis represents NA and WE reference genomes, y-axis represents the assembled contigs of NA/WE-RM-4 obtained from long read sequencing. Lower left, zoom-in dot plot of the chromosome XII inversion. The bar below represents chromosome XII, with black circle and red rectangle indicating centromere and rDNA repeats respectively. Lower right, CNV-associated with inversion structure: the extra copy of chromosome XII right arm together with the centromere was inverted and replaced the left arm. **(c)** rDNA copy number per chromosome XII (upper panel) and rDNA copy number per sample (lower panel) normalized by the ancestors for all MALs binned based on the evolved condition and the copy number of chromosome XII. Boxplot: center lines = median; boxes = interquartile range (IQR); whiskers =1.5×IQR; points = outliers beyond 1.5×IQR. **(d)** Cell doubling time (upper panel) and yield (lower panel) of RM-evolved MALs phenotyped with (left) or without (right) RM. The blue and red boxplots represent MALs with and without chromosome XII gain respectively. The *p* value was calculated to compare the cell doubling time and yield between the initial and last timepoint populations by Mann–Whitney U test.

The properties of cell growth in HU and RM evolved in different directions (Fig. 1d). Cell growth became seriously impaired by HU evolution, with the cell doubling time becoming longer (mean 2.06 h *vs*. 2.42 h, *p* = 6.80×10^-7^) and the cell yield decreasing (0.82 *vs*.-0.24, *p* = 1.15×10^-3^), indicating the accumulation of strongly deleterious mutations. This fitness loss was strongest in the earliest stages of evolution (mean cell doubling time 2.06 h initial *vs*. 2.23 at 15 single-cell bottleneck, *p* = 0.009) and then slowed substantially (mean cell doubling time = 2.23 h vs. 2.42 h at bottlenecks 15 and 60, *p* = 0.059). In contrast, no fitness loss was observed during RM evolution, with the cell doubling time remaining unchanged (mean = 1.99 h vs. 2.03 h, *p* = 0.16) and the cell yield actually improving (mean = 0.15 *vs*. 0.46, *p* = 1.18×10^-4^). These negative or neutral fitness trajectories for cell populations evolving across random single cell passages stand in stark contrast with the strongly positive fitness trajectories of the same lineages when evolving in large cell population sizes under HU and RM exposure^8^ (mean cell doubling time of adapted clones appearing after ~50 generations = 1.91 h and 1.48 h in HU and RM respectively). While our MALs experienced no selection for faster or more efficient growth, the cell viability increased somewhat for MALs evolving in YPD and HU (Fig. 1b, 1.37-fold and 1.75-fold increase), reflecting the unavoidable selection against lethal mutations during the single-cell bottlenecks. Both HU and RM evolved MALs lost some of their capacity to grow in the drug-free condition (Fig. 1c, Supplementary Fig. 3), which again is consistent with the accumulation of deleterious mutations. Taken together, the single cell passages resulted in a largely neutral evolution and the accumulation of deleterious mutations that impaired cellular fitness at different magnitudes.

### Recurrent chromosome XII amplification under RM treatment maintains rDNA homeostasis

RM inhibits cell growth by restricting the activity of the cellular master regulator TOR (Target of Rapamycin), but whether this restriction has mutagenic effects *per se* is unclear. We sequenced the MALs evolving in RM and compared the mutation spectrum to that of the YPD condition. We first focused on whole chromosome and large segmental copy number variants and structural variants. Ten out of 40 MALs (25%) evolved in YPD had at least one whole-chromosome gain or loss (Fig. 2a). This corresponded to 1.38×10^-4^ events/MAL/generation, which is similar to previous reports on the S288C lab strain evolved in YPD^10,11^ (*p*>0.16). No significant difference in the aneuploidy rate was observed among the genetic backgrounds. Among the ten MALs with aneuploidies, five acquired one extra chromosome, three acquired two or more different chromosomes, one acquired two extra chromosome I (tetrasomy) and one lost a chromosome I (monosomy). The WE/WE-YPD-4 accumulated four whole chromosomal copy number changes, which were unlikely to occur independently (*p*=5.94×10^-4^) and therefore represent connected mutational events^24^. The NA/WE-YPD-4 halved its ploidy and became a euploid haploid with recombined chromosomes (Supplementary Fig. 8), reflecting that it passed through meiosis. Globally, chromosome gains (13 events) greatly outnumbered losses (2 events) and were highly enriched in smaller chromosomes, agreeing with previous findings^10,11,14,25^. Assuming gain and loss rates to be similar, for each chromosome, this bias reflects selection against deleterious large chromosome aneuploidies and hemizygosity during evolution.

In RM, the rate of aneuploidies (4.21×10^-4^ events/line/generation) was >2-fold higher than in YPD, when excluding the unfit NA/WA-RM-4 line that underwent whole genome duplication and subsequent multiple chromosome losses and re-synthesis^26^ (Fig. 2a, Supplementary Figs. 1, 2 and 4). A highly disproportionate fraction (64.7% in RM vs 6.7% in YPD, *p*=9.54×10^-4^) of these aneuploidies corresponded to chromosome XII gains. Excluding these chromosome XII duplications, RM treatment did not alter aneuploidy rates (1.55×10^-4^ in RM *vs*. 1.38×10^-4^ in YPD, *p*=0.89). We also observed amplifications of the chromosome XII right arm in NA/WE-RM-4, spanning the rDNA-array locus and associating to an inversion of the centromeric region (Fig. 2b). RM has been reported to repress rDNA repeat amplification and lead to contraction of the rDNA repeat cluster^27^. We therefore hypothesized that complete or partial chromosome XII aneuploidies could be compensatory adaptations that counteract deleterious repeat contractions of the rDNA locus. To test this, we estimated the rDNA copy number for all the MALs from sequencing read coverage. In YPD, the rDNA copy number per chromosome XII remained unchanged (Fig. 2c, higher panel). In contrast, the rDNA copy number per chromosome XII decreased in both HU and RM conditions (*p*<0.001), consistent with loss of rDNA repeats in stress. Moreover, MALs with chromosome XII duplications had fewer rDNA repeats per chromosome XII than those with normal chromosome XII copy numbers (0.70 vs. 0.87 in HU, 0.66 vs. 0.76 in RM). Taken together, the above results support that duplication of the entire chromosome XII and of the rDNA repeat containing segments of chromosome XII are compensatory adaptations that balance RM, and possibly HU, induced rDNA repeat contraction (Fig. 2c, lower panel).

However, we found the cell doubling time evolution in RM to be unaffected by chromosome XII copy number while the cell yield gains in RM was marginally better for cells with chromosome XII gain (Fig. 2d). All the RM-evolved MALs bear a fitness cost in drug-free condition independent of chromosome XII copy number. We therefore conclude that chromosome XII gain is not adaptive in RM *per se*, which is also underscored by that the adaptive RM evolution of large cell populations of the very same lineages was driven by *TOR1* and *TOR2*^8^ mutations rather than chromosome XII duplication. Instead, chr XII amplifications are beneficial and favoured by selection only if the rDNA locus has contracted due to RM exposures, and then reverses the contraction induced fitness loss in such cells. We also confirmed that the chromosome XII duplication in RM did not destabilize the genome, because the rates of other types of mutations, i.e., substitutions, INDELs and LOH, all remained unaffected (*p*>0.23). This argues against the alternative scenario that chromosome XII duplication destabilized the genome and caused the rDNA repeat contractions.

### Chronic exposure to HU broadly elevates CNV rates

HU induced DNA replication stress is widely accepted to lead to rampant genome instability^28,29^, but the mutation rates and spectra have remained difficult to quantify precisely. We consistently observed a dramatically increased chromosome copy number variation (1.52×10^-3^ events/line/generation) in HU evolved than in RM and YPD (*p*<2.0×10^-6^) evolved cells. These included large chromosomes, loss of small chromosomes and multi-chromosome events (Fig. 2a). The two shortest chromosomes had the most frequent copy number changes (7 chromosome I losses and 7 chromosome VI gains, Supplementary Fig. 5), both being significantly more frequent than in YPD (*p*<0.021). Sixty-two percent of MALs in HU carried multiple chromosome gain/loss, greatly outnumbering those in YPD (7.7%, *p*=4.83×10^-6^). More chromosome CNV was associated with slower growth in HU (Supplementary Fig. 6a, *p*=0.017), suggesting that accumulation of aneuploidies accounts for much of the fitness loss under HU-induced mutational meltdown. Chromosome loss was associated to slower growth than chromosome gain (mean cell doubling time increase of MALs with only chromosome losses *vs*. only gains: 0.65 h vs. 0.25 h). Selection against near lethal effects of some chromosome losses likely explains why the observed rate of chromosome loss is lower than that of chromosome gain (0.45×10^-3^ vs. 1.01×10^-3^).

In addition to the whole chromosome aneuploidies, HU evolved cells often carried large segmental duplications or deletions (Fig. 3a-b) that often associated with complex structural variation. To resolve these rearranged karyotypes, we applied long-read Nanopore sequencing to two selected MALs (SA/WE-HU-2 and SA/WE-HU-3) (STAR METHODS). We found that a large-scale segmental amplification in the SA/WE-HU-2, in which an extra copy of ~84 kb SA subgenome segment at chromosome XI left end was duplicated to SA chromosome XIV right end and had replaced ~150 kb at the right arm (Fig. 3b-c). Such terminal CNV-associated translocation is usually generated by ectopic recombination or break-induced replication under replication stress^14,30^. The break points of these events coincided with tRNA and Ty elements, consistent with findings from previous genome instability studies of the *S. cerevisiae* deletion collection^31^ and the role of Ty in mediating translocations^32^. We also identified additional insertions and deletions of substantial size (mean length of insertion: 4913 bp, deletion: 322 bp), consistent with rampant genomic instability. All of them occurred within repeat sequences, including tRNA, Ty elements, X element and Y’ in subtelomere as well as within one gene with intragenic repeats (*FLO11*). Such events were more frequent in HU than in RM (Fig. 3d), suggesting that the mutagenic effects of HU also encompass these classes of structural variants.

**Figure 3.**
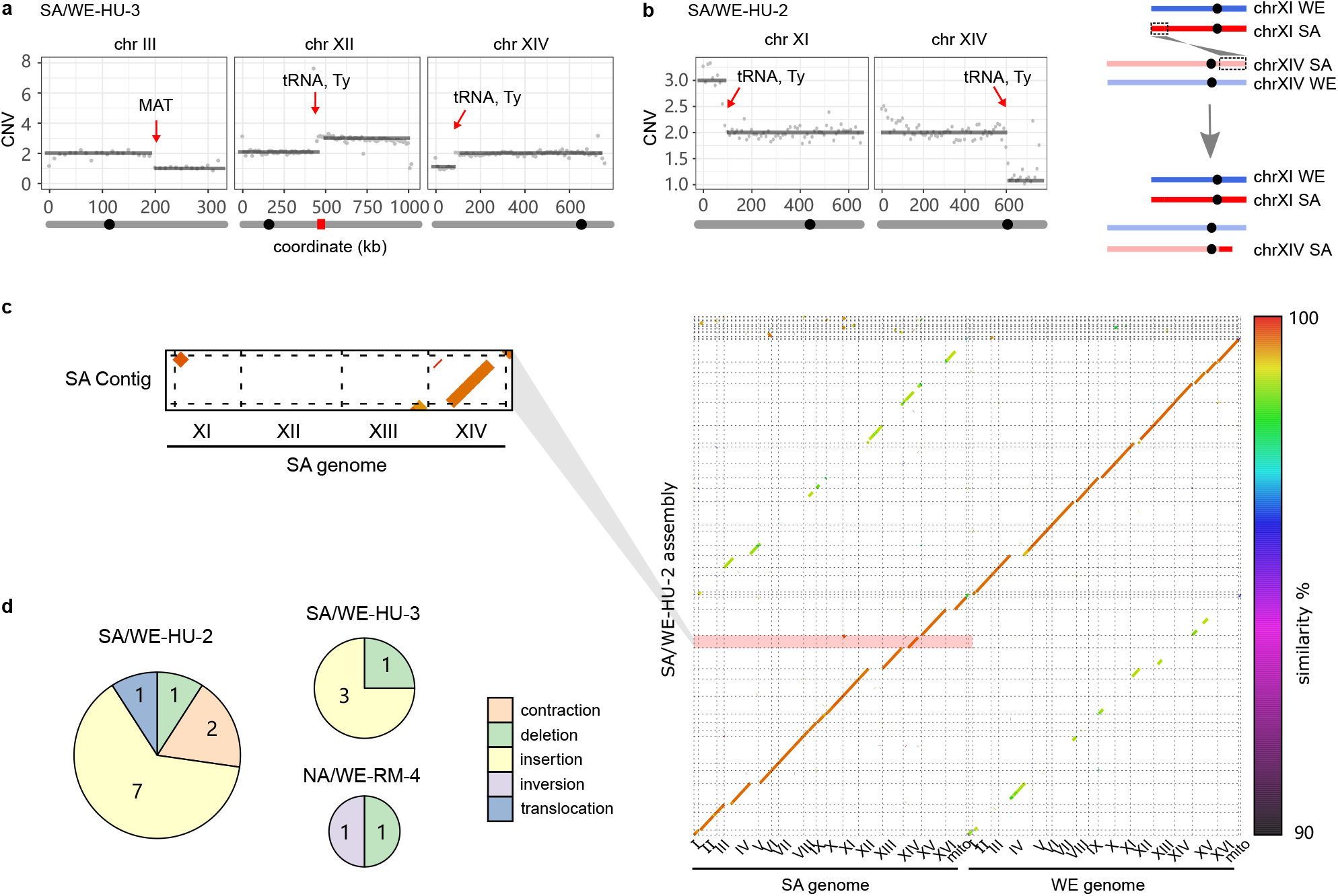
Segmental CNV and structural variation. **(a)** Segmental CNV on chromosome III, XII and XIV of the SA/WE-HU-3 line detected by short read sequencing. **(b)** Left panel, CNV on chromosome XI and XIV of SA/WE-HU-2 line detected by short read sequencing. Right panel, the gained segment on chromosome XI right end was inserted to chromosome XIV left end. In panels (a) and (b), the grey bar below the coordinate represents the chromosome, on which the black circle and red rectangle indicate the location of centromere and rDNA respectively. The red arrows indicate the segmental CNV break points. **(c)** Right panel, *de novo* phased assembly of SA/WE-HU-2 line based on the long-read sequences with color to indicate the sequence similarity between the two parental subgenomes. Left panel, the zoomed-in dot plots highlight one single assembled contig with SA chromosome XI segment inserted in chromosome XIV. **(d)** Number of structural variants identified by assembly-based alignment in three long-read sequenced MALs. Circle sizes are proportional to SVs observed and sectors indicate SVs type.

### Substitution rates remain unchanged in rapamycin but are broadly elevated in HU

We next characterized single nucleotide substitutions (hereafter substitutions) and small insertion-deletion (INDEL) in YPD, RM and HU conditions. The average substitution and INDEL rates in YPD is 1.89×10^-10^ and 1.41×10^-11^ per base per generation respectively (Fig. 4a and 4c, Supplementary Fig. 7, Tables S5-S6), consistent with previous estimates^9–12,20^. The multiple genetic backgrounds used allow us to investigate how mutation rate varies and whether background-specific mutational signatures exist. We focused on YPD condition where none of the ten genetic backgrounds went extinct and combined results from MALs sharing a specific background (Fig. 4b). We found that homozygous and heterozygous WE strains had significantly higher substitution rate (Fig. 4b,*p*<0.05). The proportion of observed *vs*. expected non-synonymous substitutions (74% *vs*. 76%,*p*=0.59) and genic substitutions (72% *vs*. 74%, *p*=0.62) across all genetic backgrounds supports a close-to-neutral substitution accumulation scenario in YPD. We also measured a transition/transversion (Ts/Tv) ratio of 0.93 (expectation with no transition bias = 0.5), which is nearly identical to the previous estimates^10^ and consistent with the reported general substitution bias towards transitions in yeast^9,11,20^. Strong C-to-T transition (30.0% of substitutions) and C-to-A transversion biases (32.4% of substitutions) drove a general GC-to-AT bias (Fig. 4d, GC-to-AT/AT-to-GC mean = 2.58).

**Figure 4.**
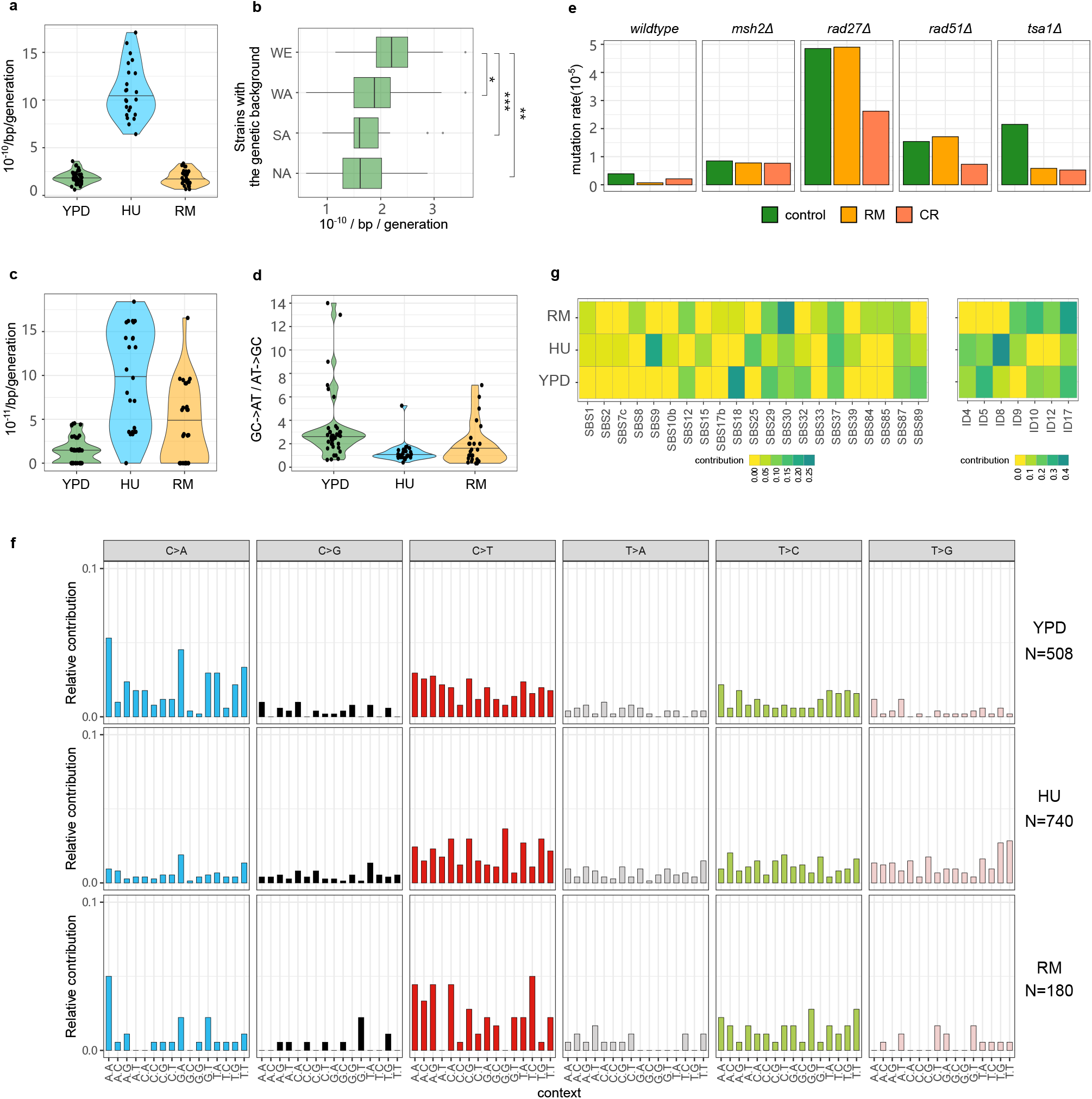
Substitutions, INDEL rates and signatures. **(a)** Substitutions rate in YPD, HU and RM. **(b)** substitutions rate of YPD-evolved MALs partitioned by the four parental genetic backgrounds, e.g., boxplot at “WE” represents substitutions rates of a total of 16 MALs of WE/WE, WA/WE, NA/WE and SA/WE in YPD. Mann–Whitney U test is used for significance analysis. \**p*=0.05, \*\**p*=0.01, \*\*\**p*=0.007. **(c)** INDEL rates in YPD, HU and RM. **(d)** Ratio of GC-to-AT to AT-to-GC substitutions. **(e)** Mutation rate estimated by fluctuation assay in wildtype and four additional mutators in BY genetic background with genes responsible for DNA repair or stress response deleted. Each strain had 16 replicates. CR is short for calorie restriction (0.5% glucose). **(f)** Relative contribution of COSMIC Single Base Substitution (SBS) and small insertions and deletions (ID) signatures. **(g)** Mutational profiles based on the relative incidences of base-substitution changes within a trinucleotide context. N indicates the number of substitutions in each condition.

The average substitutions rate in RM (Fig. 4a, 1.80×10^-10^ per base per generation) was comparable to that in YPD (*p*=0.46), while the INDEL rate (4.16×10^-11^) was only marginally higher (Fig. 4c, *p*=0.01). The proportion of non-synonymous and genic substitutions supported a neutral evolution also in RM (*p*>0.33). To confirm that RM is not generally mutagenic, we performed a canavanine-based fluctuation assay to estimate the loss-of-function mutation rate at the *CAN1* locus in the wildtype lab strain. This showed that the *CAN1* mutation rate was actually lower in RM (6.95×10^-7^) and calorie restriction (CR, partially mimicking RM, 2.12×10^-6^) conditions, compared to stress-free condition (3.90×10^-6^) (Fig. 4g). The reduced *CAN1* loss-of-function mutation rate suggested that RM may enhance the activity or fidelity of components of the DNA repair machinery, at least locally. To test this, we estimated the *CAN1* mutation rate across a panel of mutator strains. We found that RM failed to reduce the *CAN1* mutation rate in the absence of *MSH2* (DNA mismatch repair), *RAD27* (base excision and non-homologous end joining repair) and *RAD51* (nucleotide excision and double stranded break repair), but could do so in the absence of *TSA1* (anti-oxidation activity). This suggests that the DNA repair machinery is required for RM to lower the mutation rate at the *CAN1* locus. We found that T-to-C and C-to-T transitions were 1.37- and 1.41-fold more common in RM than in YPD, leading to a higher transition bias (Fig. 4e), which may reflect the activities of DNA repair pathways. Taken together, except for chromosome XII amplifications compensating for RM induced rDNA contractions, RM does not appear to be mutagenic and may stabilize parts of the genome via interactions with common DNA repair pathways.

The average substitutions and INDEL rates in HU were about six-fold higher (Tables S5-S6, Fig. 4a and 4c) than in YPD. The fraction of non-synonymous substitutions (69.8%) was marginally lower than expected (76.0%, *p* =0.03), which may signal mild purifying selection. The fraction of protein coding substitutions (73.4%) matched expectations of neutral evolution. The transition bias in HU was similar to that in YPD, but the proportion of T-to-G transversion was 3.26-fold higher (Fig. 4e), consistent with the HU mutagenic signature reported in *C. elegans*^33^. This mutation bias had the indirect effect of almost completely removing the general G/C-to-A/T mutational bias (Fig. 4d). We compared the mutation patterns observed in our HU evolved genomes to those registered in the Catalogue of Somatic Mutations in Cancer (COSMIC) (Fig. 4e-f). HU evolved cells exhibited a predominant signature of SBS9 (single base substitutions 9) and ID8 (small insertions and deletions 8), which in cancers is characteristic of DNA polymerase eta (Pol η) somatic hypermutation activity and repair of DNA double strand breaks by non-homologous end joining (NHEJ). This may reflect that HU exposed cells are forced to resort to lower fidelity DNA replication and repair processes, because of the HU block of the ribonucleotide reductase and the resulting depletion of reduced ribonucleotides^34^.

MALs with more substitutions grew faster in HU (Supplementary Fig. 6b), underscoring that it is the accumulation of aneuploidies and structural variation, rather than of substitutions, that accounts for the HU-induced fitness loss. This is compatible with a scenario in which a fraction of the accumulated substitutions might be favored by selection because they counteract the effects of some aneuploidies that otherwise would be lethal or near lethal. Indeed, gene ontology analysis shows that genes acquiring substitutions under HU evolution do not represent a random set of functions but are significantly enriched (*p*<0.0067) for genes encoding proteins binding to purine nucleotides, ribonucleotides and ribonucleoside triphosphates. Given that HU inhibits replication by inhibiting the ribonucleotide reductase^34^, such enrichment makes biological sense.

### HU is mutagenic to the mitochondrial genome

Given the huge impact of HU on virtually all nuclear mutation rates, we next estimated the copy number of mitochondrial DNA (mtDNA) to see whether the mutagenic effects encompass mtDNA. We calculated the sequencing depth of three near repeat-free mitochondrial genes (*ATP6, COX2, COX3*, Fig. 5a), which avoids much of the noise arising from the AT rich and highly repetitive nature of mtDNA^35^. We found that the rate of mtDNA loss or deletion is about 10-fold higher in HU (3.47×10^-4^ mtDNA loss/line/generation) than in RM (4.73×10^-5^) and YPD (3.29×10^-5^). A closer look at the noisy sequencing depth along the entire mitochondria (Fig. 5b-c) reveals partial mtDNA loss (heteroplasmy), which is also more frequent in HU condition (10 MALs in HU, 1 in RM and 2 in YPD). The mtDNA segments retained at normal copy numbers in these MALs ranged from 0.9 kb to 30.3 kb in size (~75-77 kb for intact mtDNA before evolution). Both mtDNA loss and deletions invariably led to *petite* colonies, reflecting a defect in oxidative phosphorylation. As yeast cells have been reported not to experience mtDNA loss under short term HU exposure (e.g. 3 days)^31^, our results suggest that HU either becomes directly mtDNA mutagenic, or indirectly causes mtDNA loss (e.g. by impairing processes necessary for mtDNA maintenance), upon chronic exposure. Although the HU induced mtDNA losses and deletions resemble those that drive adaptation under exposure to mitochondrial superoxide^36^, the HU induced mtDNA copy number changes did not associate with HU tolerance in terms of cell growth (Supplementary Fig. 6c). We also found the average mitochondrial substitution rate to be about 10-fold higher for cells evolving in HU than in YPD and RM (Fig. 5d). The elevated rates of mtDNA loss, deletions and point mutations likely reflect the lower fidelity of the mtDNA repair systems when the dNTP pool is depleted by HU^37^. Overall, we conclude that HU speeds the accumulation of a very broad range of mitochondrial mutations mirroring the nuclear genome meltdown.

**Figure 5.**
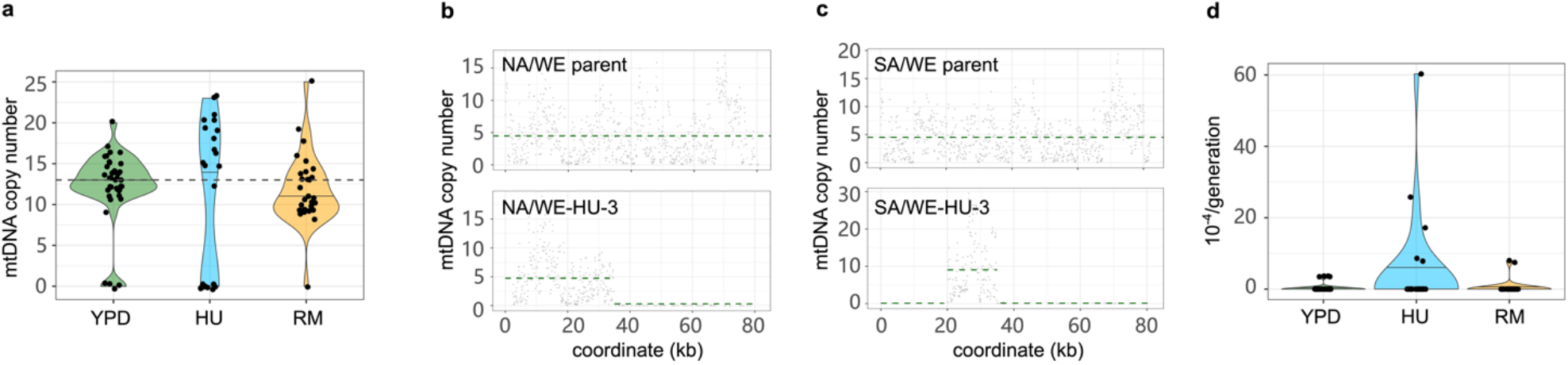
Mitochondrial DNA copy number. **(a)** Mitochondrial DNA copy number estimated by the sequencing depth of *ATP6, COX2, COX3* genes and normalized by the sequencing depth of the nuclear genome. Violin plot: center lines = median. The dashed line shows the mitochondrial copy number before evolution. **(b-c)** Mitochondrial DNA copy number of hybrid NA/WE (b) and SA/WE (c) before evolution (upper panels), and for one MAL after HU evolution (lower panels). The sequencing depth was estimated for 100 bp non-overlapping windows (dots) sliding along the mitochondrial genome (x-axis) and normalized to the sequencing depth of the nuclear genome. The dashed green line indicates the mean copy number across windows based on presence and absence. **(d)** Substitutions rate in mtDNA is higher in HU 4.86×10^-4^ substitutions/line/generation compared to 3.29× 10^-5^ in YPD and 4.73×10^-5^ in RM.

### HU increases loss of heterozygosity rates

Loss of heterozygosity (LOH) eliminates one of the parental alleles in diploid cells and allows recessive mutations in the remaining copy to affect phenotypes, which is broadly accepted to help driving cancer evolution and resistance^7,38^. We mapped LOH events in MALs of the six heterozygous strains using the MuLoYDH^12^ pipeline (Fig. 6a, Supplementary Fig. 8, Table S7) and found the overall LOH rate to be significantly higher for cells evolved in HU than in RM and YPD (Fig. 6b). Accordingly, the rates of interstitial, terminal and whole chromosome LOH in HU were 9.78-31.1 fold higher than in YPD and 4.88-7.02 fold higher than in RM (Fig. 6c). Overall, an average of 33.5% of the genome evolved in HU was affected by LOH at the end of evolution, compared with only 5.6% in YPD (*p*=9.37×10^-11^) and 6.7% in RM (*p*=2.02×10^-10^). In all environments, the average length of terminal LOH segments was much longer than that of the interstitial LOH segments and the proportion of the whole genome affected by terminal LOH was therefore 1.83~3.58 fold higher than that affected by the interstitial LOH (Fig. 6c). Such pattern is consistent with distinct DNA repair forming the LOH with interstitial events formed by two-ended homologous recombination while terminal events can be ascribed to break-induced replication^39^.

**Figure 6.**
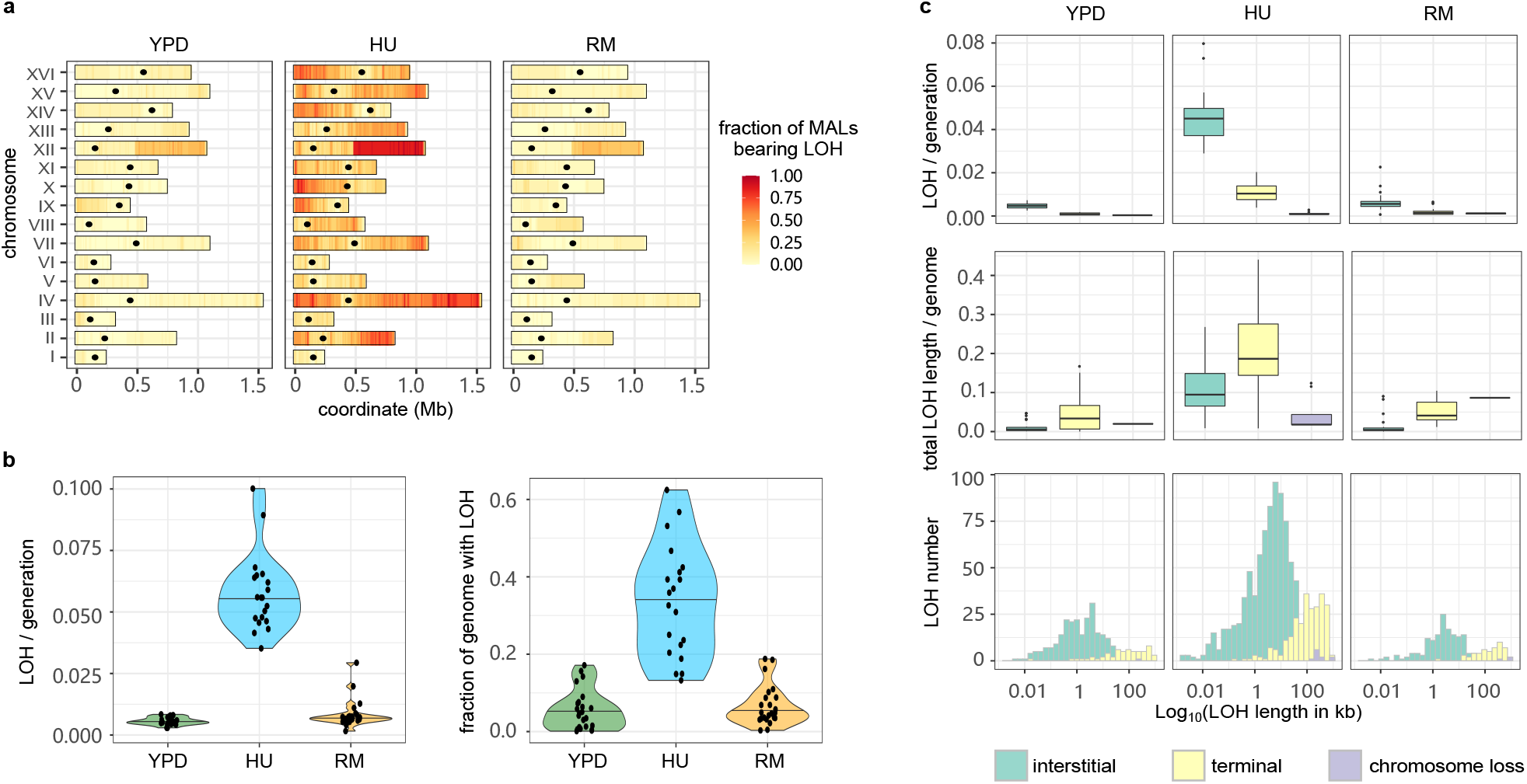
The loss of heterozygosity landscapes. **(a)** LOH fraction and genomic positions in YPD, HU and RM conditions. The heatmap shows the proportion of MALs that have LOH events using 10 kb windows across the genome. Black dots indicate the centromeres. **(b)** LOH rate (left panel) and the proportion of total LOH length in the genome (right panel). Violin plot: center lines = median. **(c)** Impact of interstitial, terminal and chromosome loss LOH type in LOH rate (upper panels), proportion of genome in LOH (middle panel) and LOH length distribution (bottom panel). Boxplot: center lines = median; boxes = interquartile range (IQR); whiskers =1.5×IQR; points = outliers beyond 1.5×IQR.

We next explored the nature of the LOH breakpoints and found their positions consistent with those previously reported in W303/YJM789 yeast hybrid evolving in a stress-free environment^14^ (Table S8). They were likely to occur in regions with high levels of γ-H2AX that marks DNA double-strand break (induced by phosphorylation of H2A histone family member X), regions with high GC content, and in non-coding RNA genes^14^. We looked closer at the breakpoints for the recurrent LOH events on chromosome XII and found them coinciding with the rDNA locus and often resulting in long terminal LOHs (Fig. 6a).

Because LOH often affects parental alleles non-randomly and drives adaptation^7,40,41^, we next examined whether parent-specific LOH bias emerged and whether the occurrence of LOH was associated with improved cell growth. We compared the LOH rate and length within each hybrid, but almost no parental biases were detected (*p*>0.06 in all conditions). The only exception was a tendency for LOH to be longer when resulting in homo- or hemizygosity for the SA background in the SA/NA hybrid evolved in YPD (*p*=0.03), where extensive homozygosity for SA associated with faster cell growth (Supplementary Fig. 9a, *p*=0.052). We also pooled the LOH events from all the hybrids involving a particular parental background to increase statistical power but found only a weak correlation between extensive homozygosity for NA and slower growth for RM evolved lineages (Supplementary Fig. 9b, *R*^2^=0.36, *p*=0.051). Thus, a parent specific LOH bias was rare among our mutation accumulation lines and had limited effects.

## Discussion

Significant efforts have been made to infer mutational signatures across the spectrum of human cancer types, therapies and environmental mutagens^42,43^, and these have provided important insights into endogenous and exogenous causes of cancer development. However, discrimination of mutagenesis from selection is difficult when working with human samples. Highly controlled conditions and single cell bottlenecks have been used to detect mutational signatures in mammalian cell lines^3,44^. This kind of approach minimized the effects of selection but mutagen treatment (e.g., chemotherapy drugs) was often short (e.g., 24 hours) and accompanied with recovery/expansion when mutagens were removed, thereby introducing a potential bias that is hard to control for. In this study, we instead applied an unconventional mutation accumulation approach by imposing continuous long-term HU and RM stresses on budding yeast. We evolved 96 yeast lines that were propagated for a total of ~181,440 generations while passing them through regular single cell bottlenecks in drug-free, HU and RM conditions, and investigated the mutational signatures. Except for an unavoidable selection against lethal mutations, which are removed in the single cell bottleneck steps, neither RM or HU imposed detectable selection that could confuse conclusions on mutation rates on evolving lines. The average spontaneous mutation rate and spectrum in drug-free condition mirrored those *S. cerevisiae* lab strains^9–11,14,20^. However, the spontaneous mutation rate varied somewhat across genetic backgrounds, showing that it is not necessarily perfectly conserved throughout a species and should be generalized with care^22^.

We revealed unique mutational signatures like T-to-G bias in HU and chromosome XII duplication in RM. RM inhibits cell growth by targeting TOR, the central signaling of cell proliferation, which is conserved from yeast to human^45^. We found the genome-wide mutation rate in RM to be comparable with that in the drug-free condition, consistent with the findings in mammalian cells that inhibition of mTOR could suppress *de novo* mutations^46^. Indeed, *CAN1*-based fluctuation assays performed in this study and by another group^47^ suggested RM may stabilize the genome locally. RM might enhance the genome stability locally by promoting DNA repair or increasing the accuracy of DNA synthesis by high fidelity polymerases. The evolution in RM also associated strongly with chromosome XII amplification, however, which did not confer growth advantage in RM. The chromosome XII amplification coincided with contraction of the rDNA repeats on this chromosome. Such RM-induced rDNA contractions are possibly near lethal, and cells can only develop into colonies to pass through single cell bottlenecks if preceded by chromosome XII amplifications that rescue the rDNA copy number. This is consistent with aneuploidy occurring at a rate that can emerge at small population sizes ranging from 1 to 10^6^ cells for each single cell bottleneck. Hence, the chromosome XII amplifications should likely not be seen as RM mutagenicity, but as an adaptive solution to the rDNA repeat contractions.

In contrast to RM, HU is highly mutagenic, resulting in an increased mutation rate in virtually all types of genomic alterations, including deleterious aneuploidies, mtDNA deletions and complex structural variations. The HU mutational signatures were compatible with cells using a low-fidelity polymerase and NHEJ for DNA replication and repair. In the absence of sex and purifying selection, such accelerated mutation rate would increase the genetic load and probably directed HU exposed populations towards mutational meltdown. As an inhibitor of ribonucleotide reductase, HU starves the cells of dNTPs and causes replication fork stalling or collapse into double strand breaks (DSBs). Frequent fork stalling can lead to un-replicated regions of DNA, which in turn cause segregation problems at anaphase^48^. This explains the observation of a high frequency of aneuploidies and segmental CNVs in HU MALs. DSBs can be repaired by cross-over or gene conversion^14,49,50^, consistent with elevated LOH in HU condition, as well as the ectopic translocation (Fig. 3b). The replication stress induced by HU also brings unique footprint in substitutions, with increased proportion of T-to-G transversions (19.9% in HU vs. 5.81% in YPD), which largely reverses the GC-to-AT bias commonly observed in yeast genome-wide mutations. This could be caused by unequal reduction of dNTP pools by HU or biased usage of dNTPs during replication stress. Further studies are needed to investigate the mechanisms responsible for such an AT-to-GC signature in HU.

In summary, we used the mutation accumulation paradigm to characterize the mutational footprint of HU and RM and drug-free conditions, findings that can be relevant to the assessment of long-term mutagenic risks of cancer therapies. The mutation accumulation approach can be extended across a wide range of different compounds and other environmental conditions, with few restrictions.

## Data availability

The genomic data generated in this study are currently being submitted and will be available under the accession code PRJNA880975.

## Acknowledgement

We thank Melania D’Angiolo, Marco Fumasoni and Sihai Yang for critical reading of the manuscript. We also thank Lorenzo Tattini for help with handling the long-read data and the MuLo-YDH pipeline. This work is supported by National Natural Science Foundation of China (32000395 to J. L. and 32070592 to J.-X. Y.), Natural Science Foundation of Guangdong Province (2022A1515011873 to J. L. and 2022A1515010717 to J.- X. Y.), Guangzhou Municipal Science and Technology Bureau (202102020938 to J. L.), Guangdong Basic and Applied Basic Research Foundation (2019A1515110762 to J.-X. Y.), Guangdong Pearl River Talents Program (2021QN02Y168 to J. L., 2019QN01Y183 to J.-X. Y.), Agence Nationale de la Recherche (ANR-15-IDEX-01), Fondation pour la Recherche Médicale (EQU202003010413), Association pour la Recherche sur le Cancer (ARCPJA32020070002320) to G. L.

## STAR METHODS

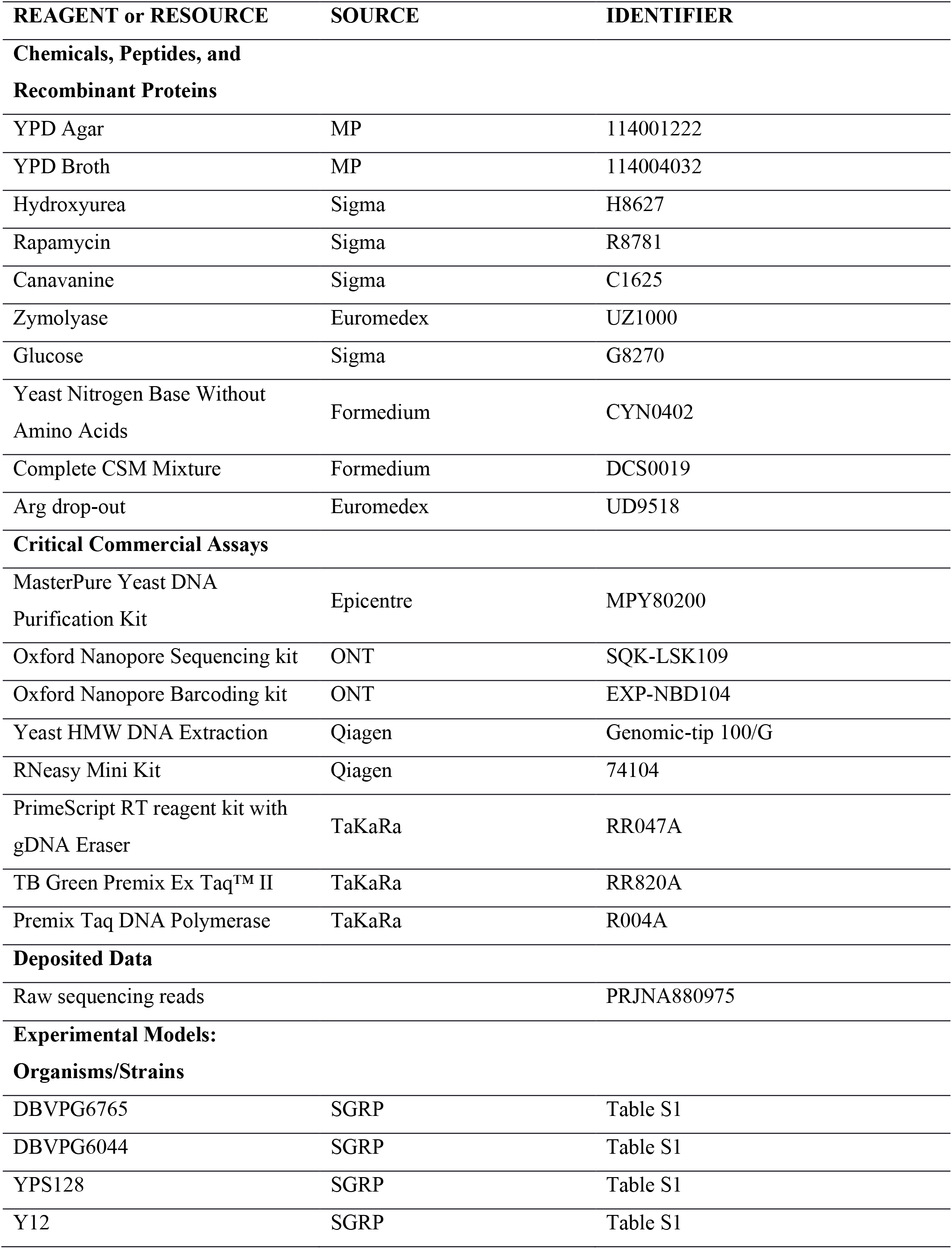
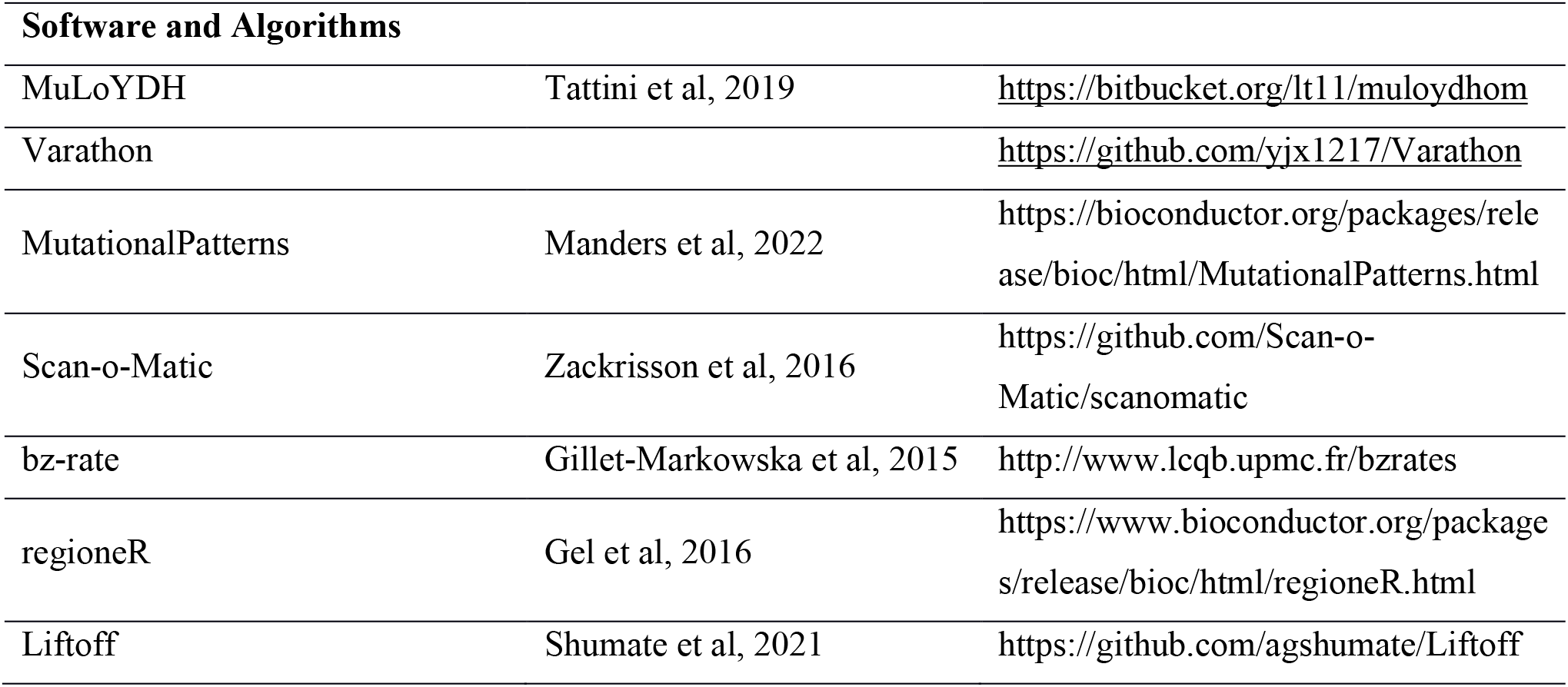

### METHOD DETAILS

#### Mutation accumulation experiments and genome sequencing

The details of strains and all the mutation accumulation lines are listed in Table S1. Briefly, ten diploid strains used in this study are derived from DBVPG6044 (West African, WA), DBVPG6765 (Wine/European, WE), YPS128 (North American, NA), and Y12 (Sake, SA), either in homozygous or heterozygous state. We propagated mutation accumulation lines (MALs) in YPD (2% peptone, 1% yeast extract, 2% glucose, 2% agar), hydroxyurea (HU, 10 mg/ml) and rapamycin (RM, 0.025 ug/ml) conditions by streaking to single colonies at fixed time intervals (2 days for YPD and 4 days for HU and RM, respectively) and letting them grow at 30 °C in a static incubator. We propagated four independent replicates for each genetic background. All the genetic backgrounds were used for MALs in YPD while two of them (WA/WA, WE/WE) were excluded in HU and RM due to low viability and severe growth defects. The Sake (SA/SA) homozygotes and SA/WA hybrids died out during mutation accumulation in HU. Therefore, there are a total of 40, 24 and 32 MALs in YPD, HU and RM, that were propagated for 120, 60 and 60 single-cell bottlenecks, respectively. We stored all the samples at five intermediate timepoints across the experiments as fossil records (single-cell bottleneck 0, 30, 60, 90, 120 in YPD; single-cell bottleneck 0, 15, 30, 45, 60 in HU and RM).

We estimated the number of generations between bottlenecks by counting the cell number (N) of the colonies using the hemocytometer. We assume that a single colony is derived from a single cell by continuous dividing without cell death. The generations between bottlenecks (G) equal log_2_(N). We estimated G for all the ancestors at P0 (G_P0_) in YPD, HU and RM conditions and for MALs at the last time point in the conditions in which they evolved (G_P60_ in HU or RM; G_P120_ in YPD). We used the mean value of G at P0 and the last time point to represent the number of generations between bottlenecks across the mutation accumulation experiments. Overall, we estimated 23 generations every 2 days for YPD, 20 generations every 4 days for HU, 22 generations every 4 days for RM, that accounted for a total of 2760, 1200 and 1320 generations across the MAL experiment for YPD, HU and RM, respectively.

The genomic DNA of all the ancestors and clones at the last timepoint was extracted with the “Yeast MasterPure” kit (Epicentre, USA) according to the manufacturer’s instructions. All these samples were sequenced by Illumina technology at Ginkgo Bioworks (Boston, Massachusetts, United States).

#### Genomic data analysis

We applied an established pipeline “MuLoYDH”^12^ that is designed for accurately tracking the mutational landscape of yeast diploids. The scripts for read mapping, coverage calculation, CNV calling and annotation, substitutions and INDEL calling and annotation, LOH detection and annotation are all embedded in MuLoYDH, which can be downloaded via https://bitbucket.org/lt11/muloydhom. For substitutions annotation we used the following criteria: i) All the substitutions must be called by both Freebayes and Samtools; ii) The quality score of each substitutions must be higher than 50 and sequencing depth at that locus must be higher than 10X; iii) Each substitutions must be supported by at least 6 reads including both forward and reverse reads. Repeat sequences such as telomeric regions were excluded. Moreover, substitutions existing in the ancestral genomes were filtered out. All substitutions and INDELs that passed these filters were visually checked in the Integrative Genomics Viewer (IGV). The ancestral genomes of WA, WE, SA and NA^51^ (https://yjx1217.github.io/Yeast_PacBio_2016/data/) were processed by the R package BSgenome (version 1.66.1, https://bioconductor.org/packages/BSgenome) to forge the corresponding reference genome packages. Then, the substitutions and INDEL mutational signatures were analyzed and fit to the COSMIC (https://cancer.sanger.ac.uk/cosmic/signatures) signatures using the R/Bioconductor MutationalPatterns package^52^, in which a strict refitting function “fit_to_signatures_strict” and a cutoff of 0.004 were applied.

In order to systematically estimate the CNV of rDNA, we applied the modules 01. Short_Read_Mapping and 04.Short_Read_CNV_Calling of the pipeline “Varathon” (https://github.com/yjx1217/Varathon). Of note, here we used a modified SGD reference which contains only one copy of the rDNA region on chromosome XII. The estimated copy number of rDNA for each MAL was normalized by that of its ancestor.

#### LOH association analysis

LOH was profiled by MuLoYDH^12^. We used “start-(first-start) ~ first” and “last ~ end+(end-last)” to define upstream and downstream breakpoints of each LOH event. We used “start ~ end” to define the LOH tracts. When analyzing the association between LOH breakpoints and published datasets and genomic features, we excluded the MALs of haploid and tetraploid. We also excluded LOH presented by only one single marker. We lift over the LOH break points as well as the Spo11 hotspots^53^ and meiotic recombination hotspots^54^ from their original reference to the SGD reference (S288C_reference_genome_R64-1-1_20110203) by Liftoff^55^. We performed permutation tests to see if the overlap between our LOH break points dataset and published datasets is higher than expected by chance using overlapPermTest function (ntimes = 1000, and alternative = “greater”) in regioneR package^56^. The output of *p* value was listed in Table S8.

#### Long-read sequencing and data analysis

We applied Oxford Nanopore Technologies (ONT) on three MALs which showed potential structural variation (SV) based on the Illumina sequencing. High-molecular weight (HMW) DNA was prepared by QIAGEN Genomic-tip 100/G. We started with 2 ug HMW DNA to prepare the sequencing library. DNA repair and end preparation were performed using the following reaction setup: 48 ul DNA, 3.5 ul NEBNext FFPE DNA Repair Buffer, 2 ul NEBNext FFPE DNA Repair Mix, 3.5 ul UltraII End-prep reaction buffer and 3 ul UltraII End-prep enzyme mix, which was incubated at 20 °C for 15 minutes followed by 65 °C for 15 minutes. After AMPure XP Beads clean-up (1:1 ratio), different native barcodes (ONT, EXP-NBD104) were ligated to each sample (22.5 ul DNA, 2.5 ul Native Barcode and 25 ul Blunt/TA ligase Master Mix, incubated at 25 °C for 20 minutes). After AMPure XP Beads clean-up (1:1 ratio), multiple barcoded samples were pooled and sequencing adaptor was ligated to the pooled library (65 ul pooled DNA, 5 ul AMII, 20 ul NEBNext Quick ligation reaction buffer, 10 ul Quick T4 DNA ligase, incubated at 25 °C for 15 minutes). Following AMPure XP Beads clean-up (0.4X) and L Fragment Buffer wash (ONT, SQK-LSK109), the library was eluted in 15 ul Elution Buffer. The library was loaded into the FLO-MIN106 MinION flow cell according to the manufacturer’s guidelines (ONT, SQK-LSK109) to perform the whole-genome long-read sequencing.

The raw nanopore reads were processed using the 00.Long_Reads module of the LRSDAY v1.5.0 framework^57^. Briefly, we performed base calling and demultiplexing with Guppy v2.3.5. The demultiplexed reads were further processed by Porechop v0.2.4 (https://github.com/rrwick/Porechop) for adaptor trimming (option: – discard_middle). The *de novo* assembly was carried out by the 01.Long-read-based_Genome_Assembly module of LRSDAY. We used “canu” as the assembler (Koren et al, 2017; https://github.com/marbl/canu). SVs were called by MUM&Co^58^, which uses Whole Genome Alignment information provided by MUMmer (v4) to detect variants. The SVs were further crossvalidated with the results called by the module 11.Long_Read_Mapping of the pipeline “Varathon” (https://github.com/yjx1217/Varathon) and manually checked in IGV.

#### Phenotyping

We revived the intermediate samples of MALs and phenotyped them by measuring their doubling time and yield. We used a high-resolution large-scale scanning platform, Scan-o-matic, to monitor cell growth in a 1536-colony design on solid agar plate^59^. The Scan-o-matic program uses data from the time-series images taken by the high-quality desktop scanners to calculate the population size and generate growth curves for the colonies. Phenotyping was run for 3 days, and scans were continuously performed every 20 minutes. The conditions of phenotyping were the same as that in the mutation accumulation experiments. Each sample had at least 6 technical replicates. After quality control filtering, the measurement of doubling time was extracted for downstream analysis in R (R version 3.6.3). Yield used in the paper was a log2 ratio between a surface of the assay position and the spatial controls. All the scripts are available on GitHub (https://github.com/Scan-o-Matic/scanomatic; last accessed September 1, 2020).

#### Viability estimation

To estimate the cell viability in drug conditions, we cultured cells in liquid YPD until they reached mid-log phase and then diluted the culture to plate ~200 cells on solid medium with or without drugs. After incubation at 30 °C for 4 days, we counted colony-forming units (CFU) and calculated the viability based on the ratio of CFU in HU/RM and YPD control conditions. To estimate the viability in YPD condition, we measured CFU on YPD plates using the same approach as described above. Each strain had four replicates. Meanwhile, we count the cells under the microscope by hemocytometer. After transforming CFU and counted cell number to the same dilution scale, viability was calculated by CFU divided by cell number.

#### Fluctuation assay

The protocols of Luria-Delbrück fluctuation assay^60^ were applied to determine the mutation rate at which the cells become resistant to canavanine (60 ug/ml) in different conditions. The strains used for this experiment were gifts from Dr. Alain G. Nicolas^15^. Briefly, 16 clones of each strain were picked, then independently diluted to ~100 cells and cultured in SC (2% glucose, 0.675% yeast nitrogen base, 0.088% complete amino acid supplement, pH=6.0), SC+RM (0,025 μg/mL) and SC with calorie restriction (0.5% instead of 2% glucose), respectively. After four days of incubation at 30 °C, we spotted the cultured cells on the over-dried canavanine plates, as well as the SC plates as control, using an appropriate dilution to avoid too many or too few cells. After three days of incubation, we counted the resistant colonies and inferred the number of mutants and total cells in the culture of SC, RM and calorie restriction. We used an online tool “bz-rate” to calculate the mutation rate for different conditions^61^ (http://www.lcqb.upmc.fr/bzrates).

## Notes

### Competing Interest Statement

The authors have declared no competing interest.

### Summary of Updates

An updated version of our previous submission with improved text and figures

