## supplementary figures for "Genome instability footprint under rapamycin and hydroxyurea treatments"

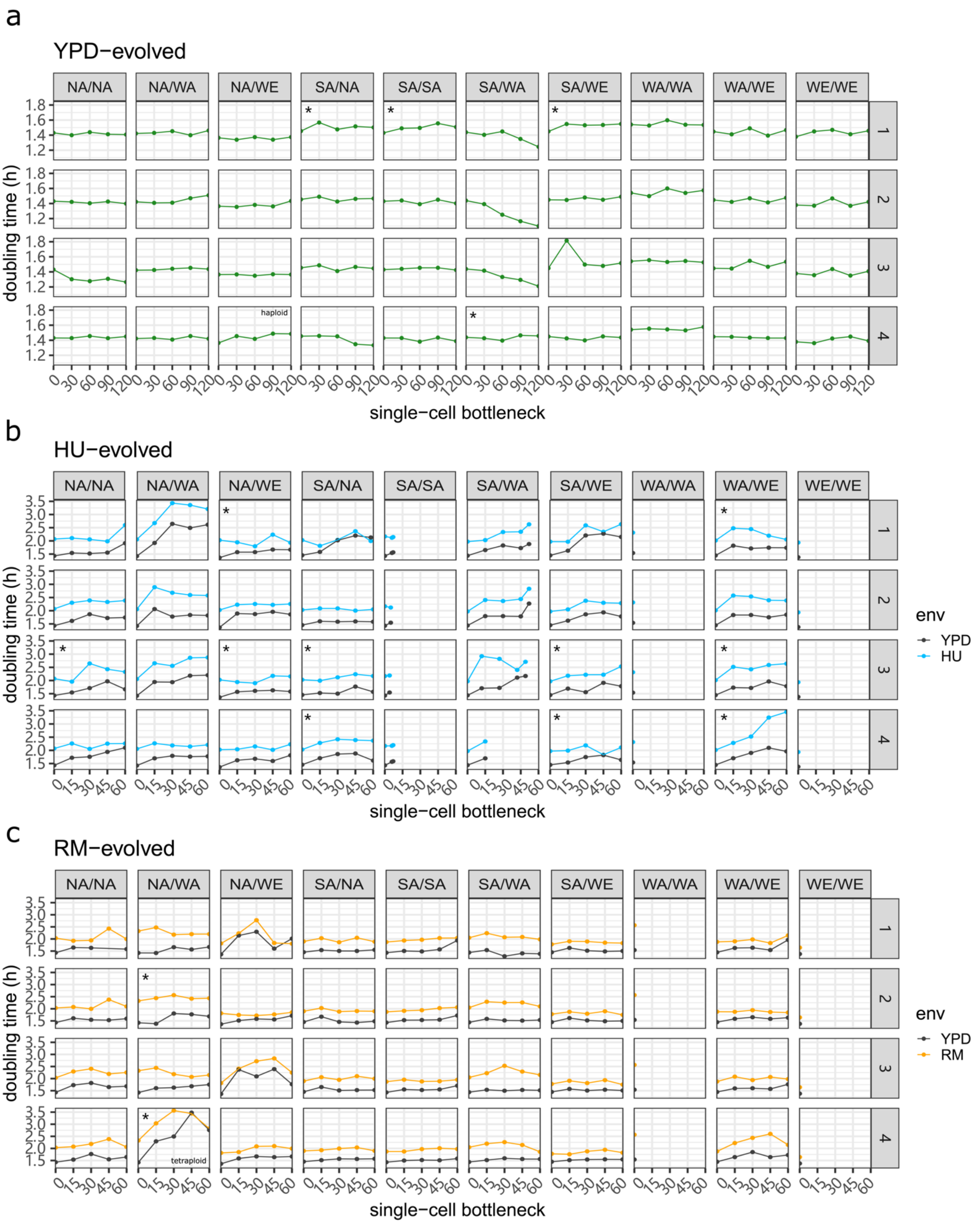


**Supplementary Figure 1.** The doubling time dynamics of the **(a)** YPD-evolved, **(b)** HU-evolved, and **(c)** RM-evolved MALs across five time points during mutation accumulation. The green, blue, and orange dots show the doubling time (hours) measured in YPD, HU and RM condition respectively while the grey dots show the doubling time of drug-evolved MALs phenotyped in drug-free condition. The NA/WA-RM-4 line revealed strong growth defects in both RM and YPD conditions, consistent with its extensive unbalanced chromosome number (mean doubling time 2.82 h *vs.* 2.26 h of other NA/WA MALs in RM condition, 2.76 h *vs.* 1.70 h in YPD condition). “*” indicates mtDNA loss.

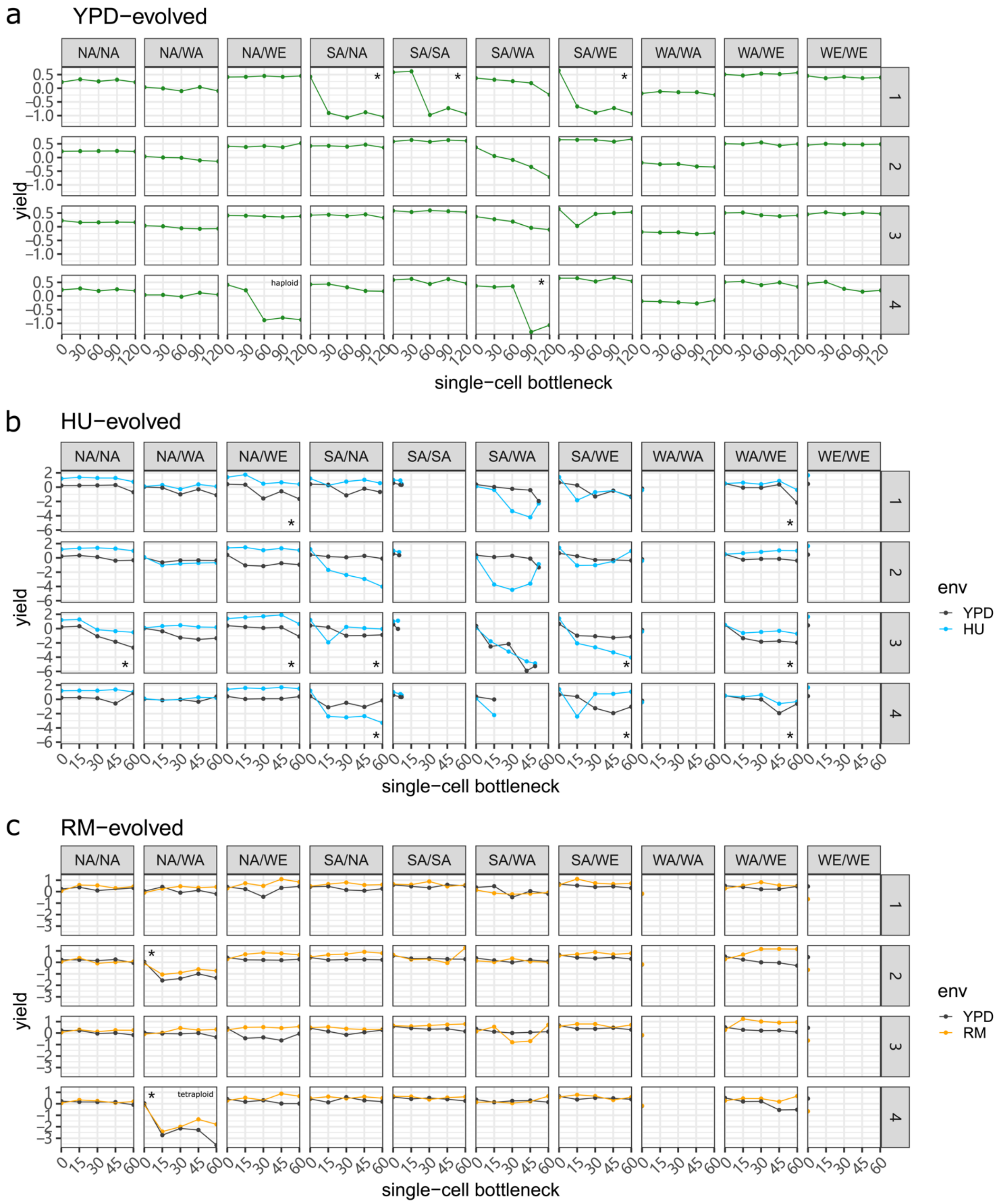


**Supplementary Figure 2.** The dynamics of relative yield of all the **(a)** YPD-evolved, **(b)** HU-evolved, and **(c)** RM-evolved MALs across five time points during mutation accumulation. The green, blue, and orange dots show the relative yield measured in YPD, HU and RM condition respectively while the grey dots show the yield of drug-evolved MALs phenotyped in drug-free condition. "*" indicates mtDNA loss.

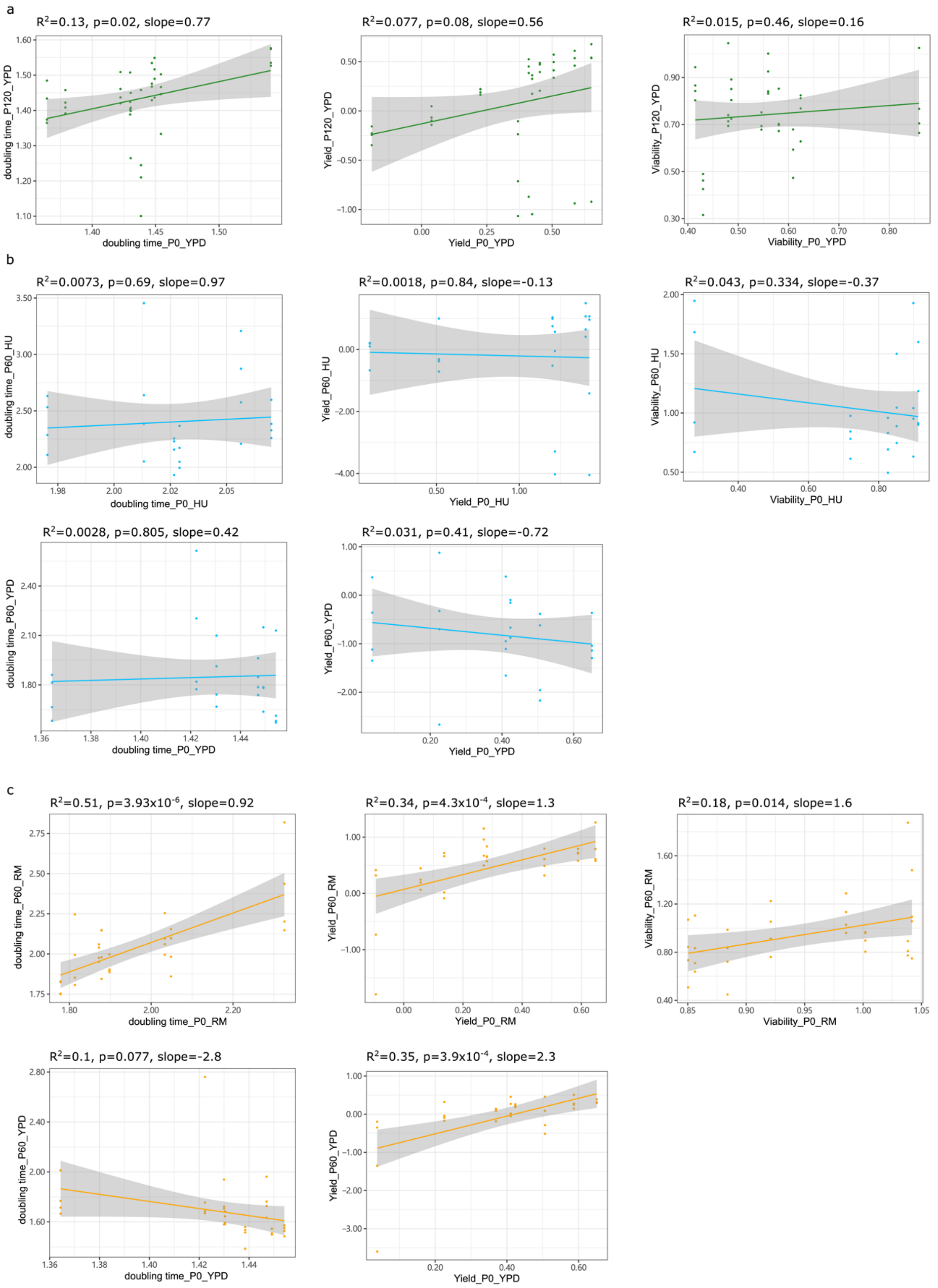


**Supplementary Figure 3.** Phenotypic correlations between different time points and different conditions for MALs evolved in **(a)** YPD, **(b)** HU and **(c)** RM. For (a-c), left panel: doubling time correlation; middle panel: yield correlation; right panel: viability correlation. For (b) and (c), from up to bottom the correlation analysis includes comparison between the initial time point and end time point phenotyped in drugs (upper panel), phenotyped without drugs (middle panel), and comparison between phenotypes with and without drugs at the end time point (bottom panel).

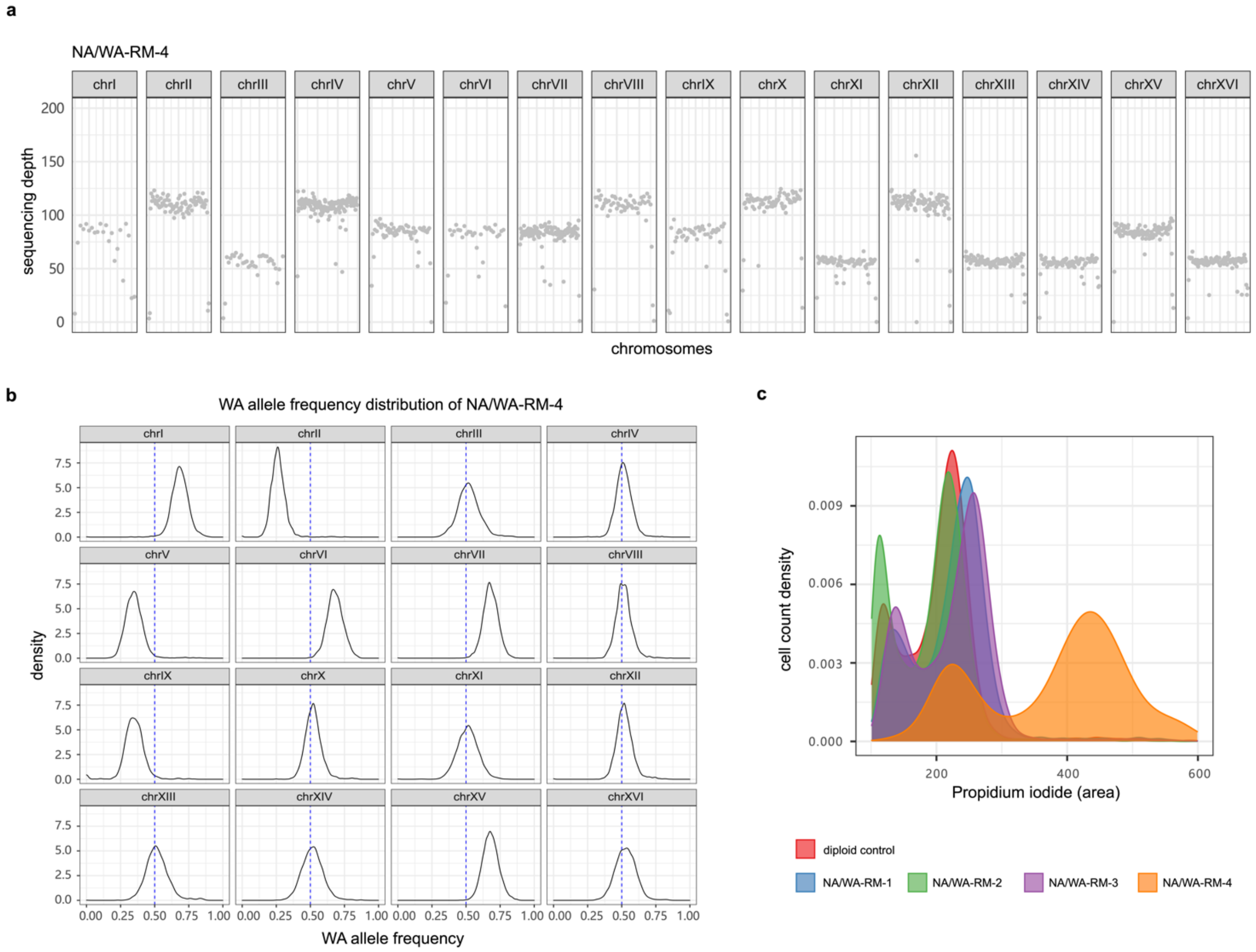


**Supplementary Figure 4.** The NA/WA-RM-4 line underwent whole genome duplication and subsequent multiple chromosome loss and re-synthesis. **(a)** Sequencing depth of NA/WA-RM-4 across all the chromosomes. **(b)** WA allele frequency distribution of NA/WA-RM-4 on each chromosome. Chromosomes II and XV are present in four copies but with an unbalanced WA allele frequencies (1:3 and 3:1 respectively). Such scenario is consistent with loos of one copy and subsequent resynthesis using the other parent homolog as template. We described a similar situation in a MAL initiated with a natural *S. cerevisiae* x *S. paradoxus* hybrid that also experienced WGD duplication and subsequent chromosome loss and re-synthesis cycles, suggesting that this mechanism is prevalent in tetraploids (D'Angiolo 2020) **(c)** Ploidy flow cytometry profiles for all the four NA/WA-RM- evolved lines.

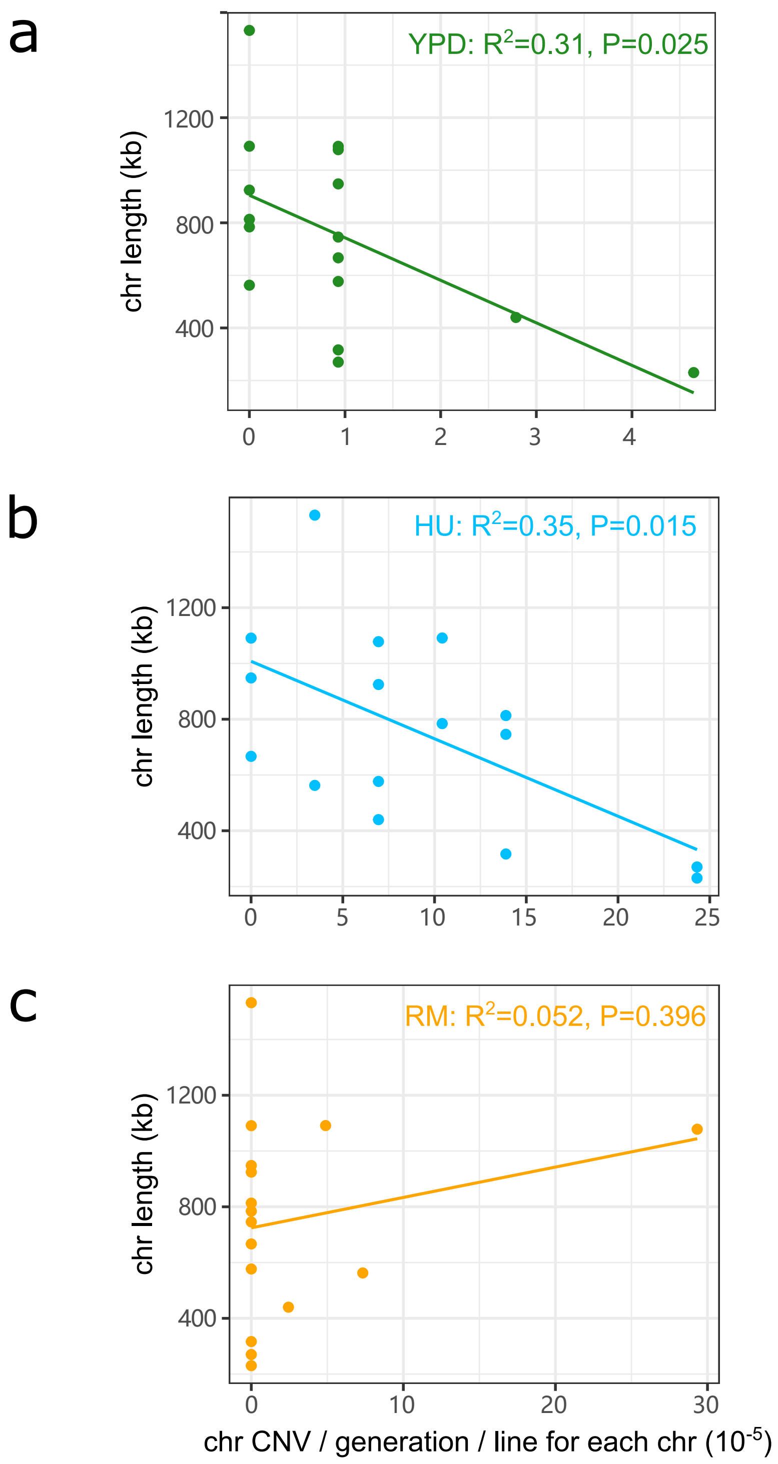


**Supplementary Figure 5.** The correlation of chromosome length and chromosomal CNV across MALs in **(a)** YPD, **(b)** HU and **(c)** RM condition.

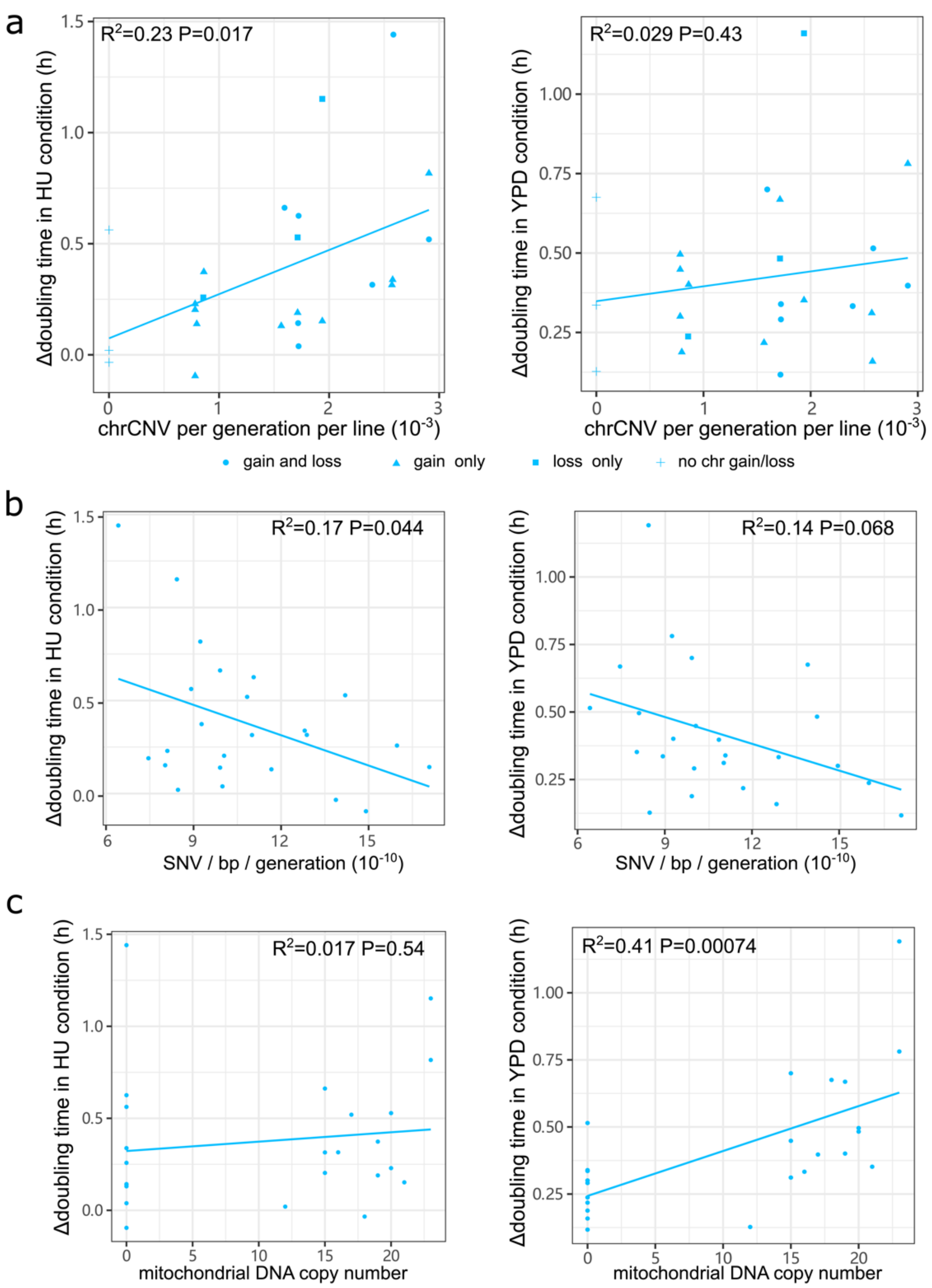


**Supplementary Figure 6. Correlation between mutations and phenotypes. (a) C**orrelation between whole chromosome CNV rate in HU and doubling time change (last time point - initial time point) measured in HU (left panel) and without HU (right panel). The triangle, square, circle and cross represent MALs with chromosome gain only, loss only, both gain and loss, no whole chromosome CNV. **(b) C**orrelation between substitution rate in HU and doubling time change (last time point - initial time point) measured in HU (left panel) and without HU (right panel). **(c) C**orrelation between mitochondrial DNA copy number in HU and doubling time change (last time point - initial time point) measured in HU (left panel) and without HU (right panel).

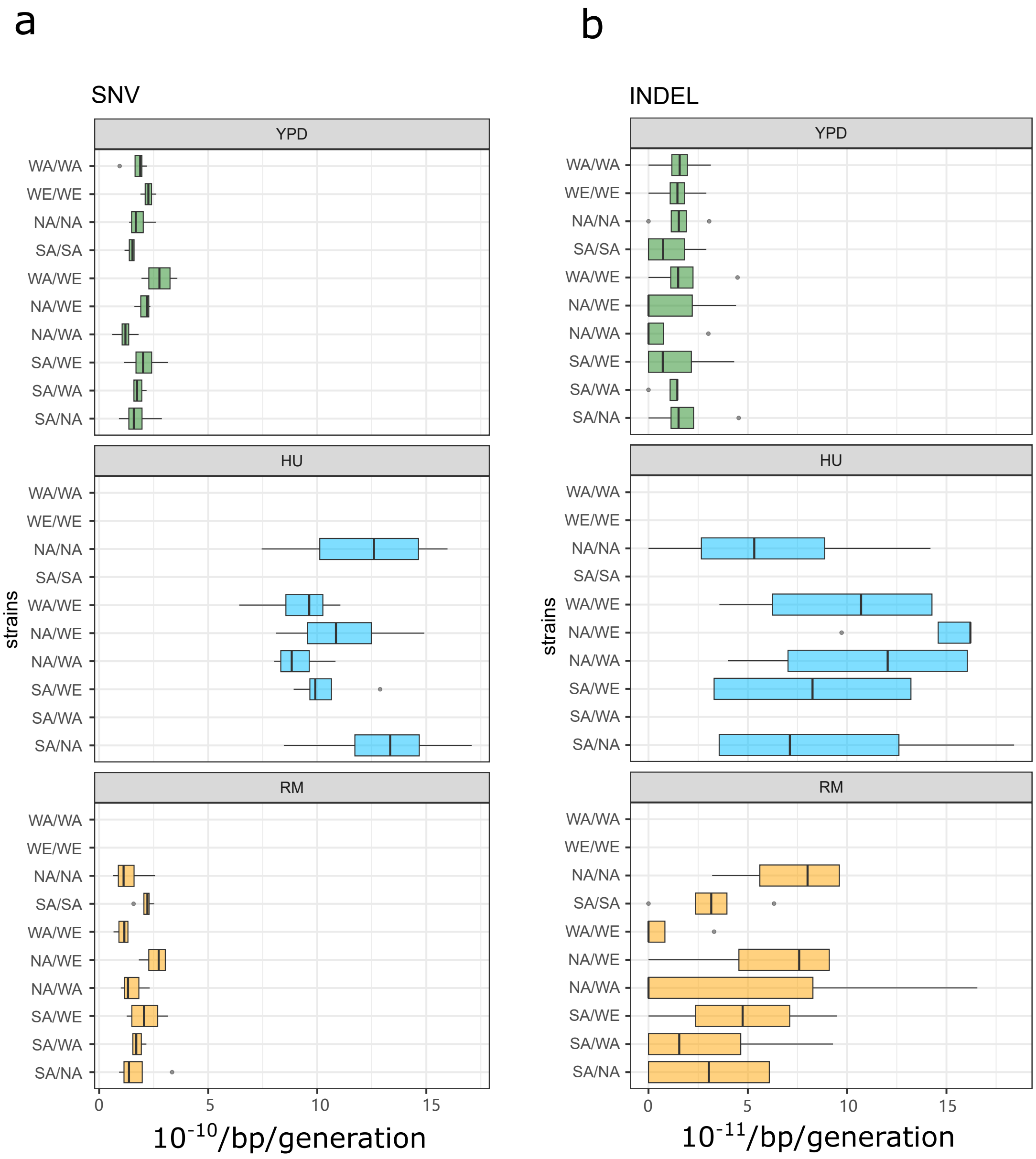


**Supplementary Figure 7.** The rate of **(a)** substitution and **(b)** INDEL for each MAL partitioned by the genetic background. Each panel from up to bottom shows the mutation rate in YPD, HU and RM condition respectively.

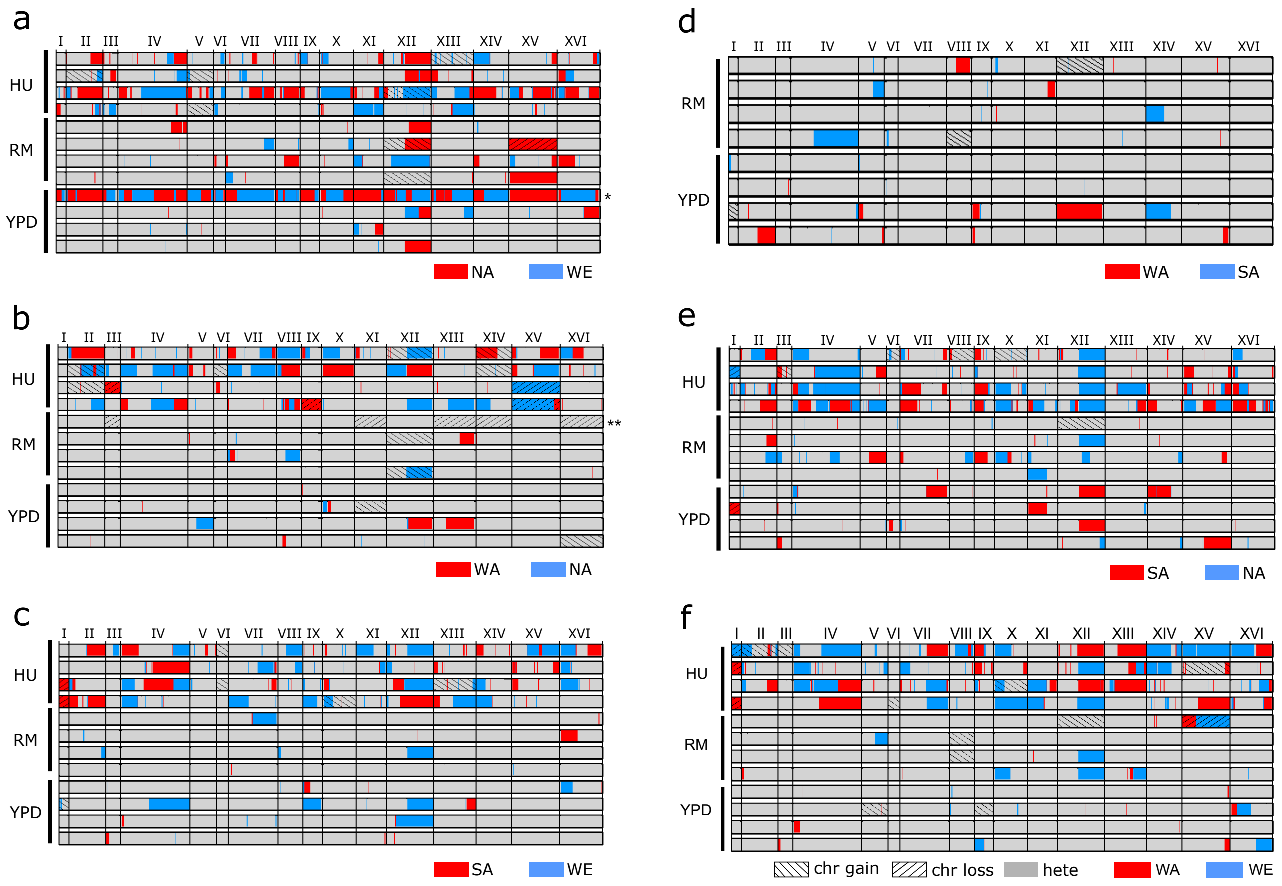


**Supplementary Figure 8.** Genome-wide LOH landscape of hybrid mutation accumulation lines (MALs) for **(a)** NA/WE, **(b)** NA/WA, **(c)** SA/WE, **(d)** SA/WA and **(e)** SA/NA **(f)** WA/WE. Each panel shows the LOH landscape of one MAL (from bottom to top represents replicates 1 to 4). The blue and red blocks represent the LOH events towards one of the parents. The grey blocks represent the heterozygous status. The blocks with slash or backslash indicate chromosome loss or gain respectively.

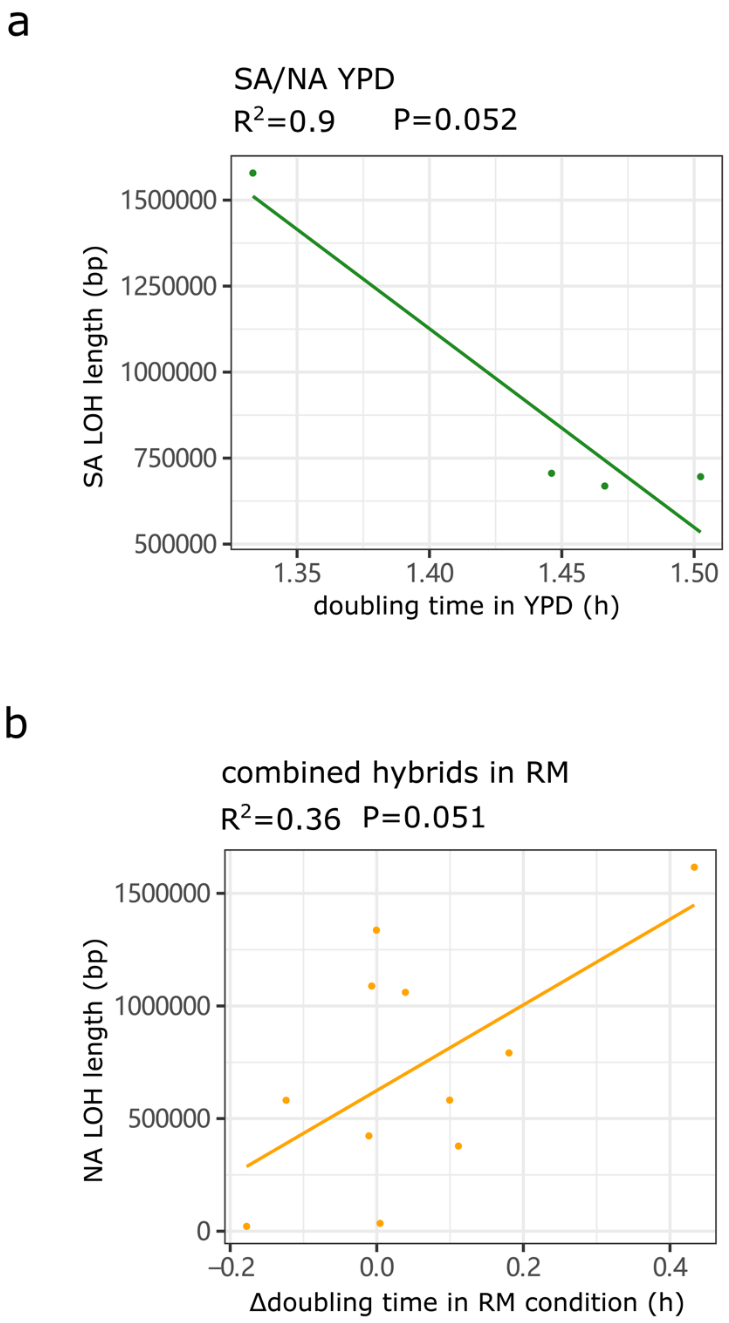


**Supplementary Figure 9. Total LOH length and its effects on fitness. (a)** Correlation of the total LOH length of SA allele of SA/NA hybrid (four replicates) and its doubling time in YPD. **(b)** Correlation of the total LOH length of NA allele of all the hybrids (11 MALs) and their doubling time change in RM.

**Supplementary tables**

**Table S1.** Strains and mutation accumulation lines

**Table S2.** Doubling time (hours) and yield of all the MALs across multiple time points during mutation accumulation

**Table S3.** Viability of the ancestors and drug-evolved mutation accumulation lines

**Table S4.** Viability of the ancestors and YPD-evolved mutation accumulation lines

**Table S5.** List of identified substitutions

**Table S6.** List of identified INDELs

**Table S7.** List of identified LOHs

**Table S8.** LOH breakpoints association with published datasets and genomic features
